# Deciphering the global genomic landscape of *C. neoformans*: Population dynamics, molecular epidemiology and genomic signatures of pathogenicity

**DOI:** 10.1101/2025.11.11.687764

**Authors:** Jananishree Sathiyamoorthy, Jayapradha Ramakrishnan

**Author notes:** Corresponding Author: Jayapradha Ramakrishnan, Actinomycetes Bioprospecting Lab, Centre for Research in Infectious Diseases (CRID), School of Chemical and Biotechnology, SASTRA Deemed University, Tirumalaisamudram, Thanjavur – 613401, Tamil Nadu, India.

## Abstract

The study investigates global genomic surveillance of *Cryptococcus neoformans* over three decades to elucidate genetic diversity, incorporating serotypes, molecular types, STs and mating types, phylogenomics, virulence-associated determinants, antifungal resistance, and pangenome profiles. Across 139 study genomes, the isolates exhibited nineteen distinct sequence types (STs), among which ST5 (63%) was the most prevalent. The predominance of MATα mating type 91% mirrors enhanced virulence, environmental adaptability and clonal expansion, enabling its persistence without MATa (9%) counterparts. The phylogenomic analysis revealed, there is no distinct clustering of isolates from clinical and environmental settings, indicating a high level of genome conservation across sources. Core–accessory genome partitioning and ortholog-based clustering revealed clonal population dominance of the VNI lineage across continents, alongside clear divergence from the VNIV lineage. Virulence gene analysis highlighted conserved expression of capsular genes, phospholipase gene, iron acquisition genes and superoxide dismutase genes, while point mutations and aneuploidy in *ERG11* and *AFR1* suggested potential azole-resistance mechanisms. This study delivers a holistic assessment of global genomic diversity in the *C. neoformans* population, emphasising evolutionary conservation, lineage-specific divergence, and adaptability to environment, host and anti-fungal treatments.

## 1. Introduction

*Cryptococcus neoformans* is an environmental encapsulated basidiomycete. They are ubiquitous and are niche-specific. *Cryptococcus* is a sapronotic infection that causes opportunistic and occasionally primary infections in humans^1,2^. Its global distribution spans tropical, subtropical and temperate regions, serving as persistent sources of infection. Globally, cryptococcal meningitis accounts for an estimated 1, 52, 000 cases reported annually and 1,12,000 deaths worldwide, leading to 74% of mortality in patients with immunocompression^3^. The global mortality profile depicts the disproportionate burden in Sub-Saharan Africa, underscoring its clinical and public health significance^4^. Among various vulnerable populations, patients with advanced HIV infection are reported as being the most susceptible to acquiring cryptococcal infection^3,4^. *C. neoformans* has diverse clinical manifestations, ranging from localized pulmonary disease to disseminated CNS infections, underscoring its pathogenic versatility.

*C. neoformans* is categorized as a critical priority pathogen, due to its high mortality rate, inherent and acquired antifungal resistance, an extensive virulence repertoire and evolutionary fitness attributed to exceptional ecological adaptability^5^. The key adaptability traits include polysaccharide capsule, melanin, urease, phospholipases and thermotolerance, all of which help the pathogen to survive in the environment and human host^6,7^.

*C. neoformans* reproduces mainly by budding (asexual), especially inside the host due to nutrient-rich conditions. Sexual reproduction occurs between compatible mating types (a*α/ α*α) and leads to basidiospore formation, but this typically happens under nutrient-limited conditions, especially nitrogen deprivation, dehydration and high cell density. In nature, asexual reproduction is relatively rare but contributes to genetic diversity by unisexual mating. Genetic diversity has revealed two subspecies (.*grubii* and .*neoformans*), three serotypes (A, D and AD) and five major molecular types (VNI, VNII, VNIII, VNB and VNIV) with evidence of both clonal expansion and recombination events shaping population dynamics^8–10^.

The advent of high-throughput sequencing has revolutionized fungal genomics, elucidating comprehensive insights into genetic diversity, population structure, evolutionary relationships and molecular framework of pathogenic adaptation^9,11^. Sequencing-based approaches enable detailed study of genetic variations, as well as gene family expansion and contraction across fungal lineages. Furthermore, genomic interrogation of virulence determinants and antifungal resistance aids in developing potential targeted therapeutics and diagnostic strategies. On the whole, the sequencing technologies have become indispensable in fungal research, supporting progress from basic research to the development of novel antifungal interventions and disease control.

Keeping this as a basis, the present study aimed to utilize a comprehensive comparative genomics to investigate evolutionary biology in *C. neoformans* over three decades (1998-2021), analyzing 139 publicly available strains from diverse environments and host factors worldwide. Considering the origin of strain, year of isolation, ST, serotypes, molecular types, mating type, resistome, and virulome for the comparative analysis. Further, the genomic diversity was characterized by pangenome analysis. To the best of our knowledge, this constitutes the first study to perform a global comparative analysis of *C. neoformans*.

## 2. Materials and Methods

### 2.1. Retrieval of assembled genome and metadata

In total, 183 genomes of *C. neoformans* from the NCBI dataset (from 1970-January 2025) were retrieved. Out of which 139 genomes were included for this study and 44 genomes were excluded due to undefined isolation sources. The metadata of the isolates were compiled from NCBI datasets and complemented with information (ST type, serotype, mating type and molecular type) retrieved from previously published studies (supplementary table 1, sheet 1).

### 2.2. Comparative whole genome diversity analysis

The EZBioCloud platform, a web-based tool, was used to compare the whole genome nucleotides, to understand their similarities, and the results were recorded in MS Excel sheet for visualisation, as there is no visualisation tool^12^.

### 2.3. Whole-genome SNP analysis

For the whole-genome SNP analysis, the assembled reads were mapped using “Map with BWA-MEM-Galaxy v0.7.19“ to the reference genome (H99)^13^. To determine the SNPs, “Freebayes-Galaxy v1.3.10+galaxy0” was applied^14^. Implementing the “VCFfilter in Galaxy v1.0.0_rc3+galaxy3,” variants with low quality scores, large error rates, or unclear calls were eliminated. The VCF file was converted into a tabular file with Galaxy v4.3+t.galaxy0’s SnpSift Extract Fields.

### 2.4. Phylogenomic Delineation

The evolutionary relationship among the 139 genomes was assessed using OrthoFinder (Galaxy Version 2.5.5+galaxy1), where orthologous genes were used and the tree was constructed based on the maximum likelihood method^15^. In order to investigate the phylogenetic placement based on MLST, the seven housekeeping genes (*CAP59*, *GPD1*, *PLB1*, *SOD1*, *LAC1*, *IGS1* and *URA5*) were concatenated for each genome and a reference strain, H99. Using the MegaX platform maximum likelihood method was used to view the tree with a bootstrap value of 1000. Tree editing and annotation were performed using the interactive Tree of Life (iTOL)^16,17^.

### 2.5. Pan-genome analysis

The pan-genome analysis was executed with OrthoFinder (Galaxy Version 2.5.5+galaxy1)^15^. The output from Orthofinder was visualized using iTOL for phylogenetic tree construction and the Linux platform (matplotlib, plotly, pandas and Seaborn) for core and accessory gene visualizations^18,19^.

### 2.6. Virulome Diversity Analysis

The major virulence genes were collected from the literature and the DFVF database^20–22^. The protein sequences of each major virulence gene (n=17) were retrieved from Uniprot and their presence in the annotated genomes was identified using pBLAST^23,24^. The sequence with 99 to 100% identity was considered for the presence of VRGs. The comprehensive result of the 17 major virulence genes was consolidated using iTOL.

### 2.7. Resistome Diversity Analysis

The mutations responsible for the development of Anti-Fungal Resistance (AFR) provided in the AFRbase database were compared with the annotated genomes of the strains using BLAST and CLUSTAL Omega^25^. The results of the comparative AFR and mutation profiles were consolidated using iTOL. Following, the mutation analysis was also performed using the ChroQuesTes tool to check the manually interpreted results^26^. Further, the results were analysed by *in vitro* methods, Minimum Inhibitory Concentration (MIC) and Antifungal Susceptibility testing (AFST) test using the available antifungals (Amphotericin B, Fluconazole and 5-fluorocytosine for MIC, Voriconazole, Itraconazole, Miconazole and Ketoconazole for AFST) and strains (H99 and Cn).

## 3. Results

### 3.1. Global Distribution and Epidemiology

*C. neoformans* genomes (n=139) were retrieved from the NCBI dataset. The metadata, which includes geographic location, source, ST, serotypes, molecular type, mating type locus, genome size, GC content, sequencing details, host, and associated diseases, was provided in Supplementary Table 1. The strains were categorised into 126 clinical and 13 environmental isolates. Clinical strains were primarily from cerebrospinal fluid (CSF) (n = 84, 60.43%), followed by blood (n = 13, 9.4%), bronchoalveolar lavage fluid (n = 3, 2.2%), lung tissue (n = 8, 5.8%), and skin (n=1, 0.7%). The sources of isolation establish the fact that the spectrum of cryptococcal disease ranges from self-limiting cutaneous infections to fatal systemic infections. Environmental strains were sourced from pigeon guano (n = 7, 54%), cockatoo excrement (n = 1, 7.7%), mopane tree (n = 3, 23%), and eucalyptus tree (n = 2, 15.4%). The predominance of CSF isolates highlights cryptococcal meningitis as the primary clinical manifestation. Approximately 89% of strains originated from tropical (Columbia, Brazil, Botswana, India, Thailand and Vietnam) and subtropical regions (U.S, China, Taiwan and Melborne), while 11% originated from temperate regions (U.S, France and Tanzania), indicating *C. neoformans’* potential to grow in high environmental temperatures (41 °C) and its adaptability for its survival in human hosts (Figure 1a and 1b).

**Figure 1:**
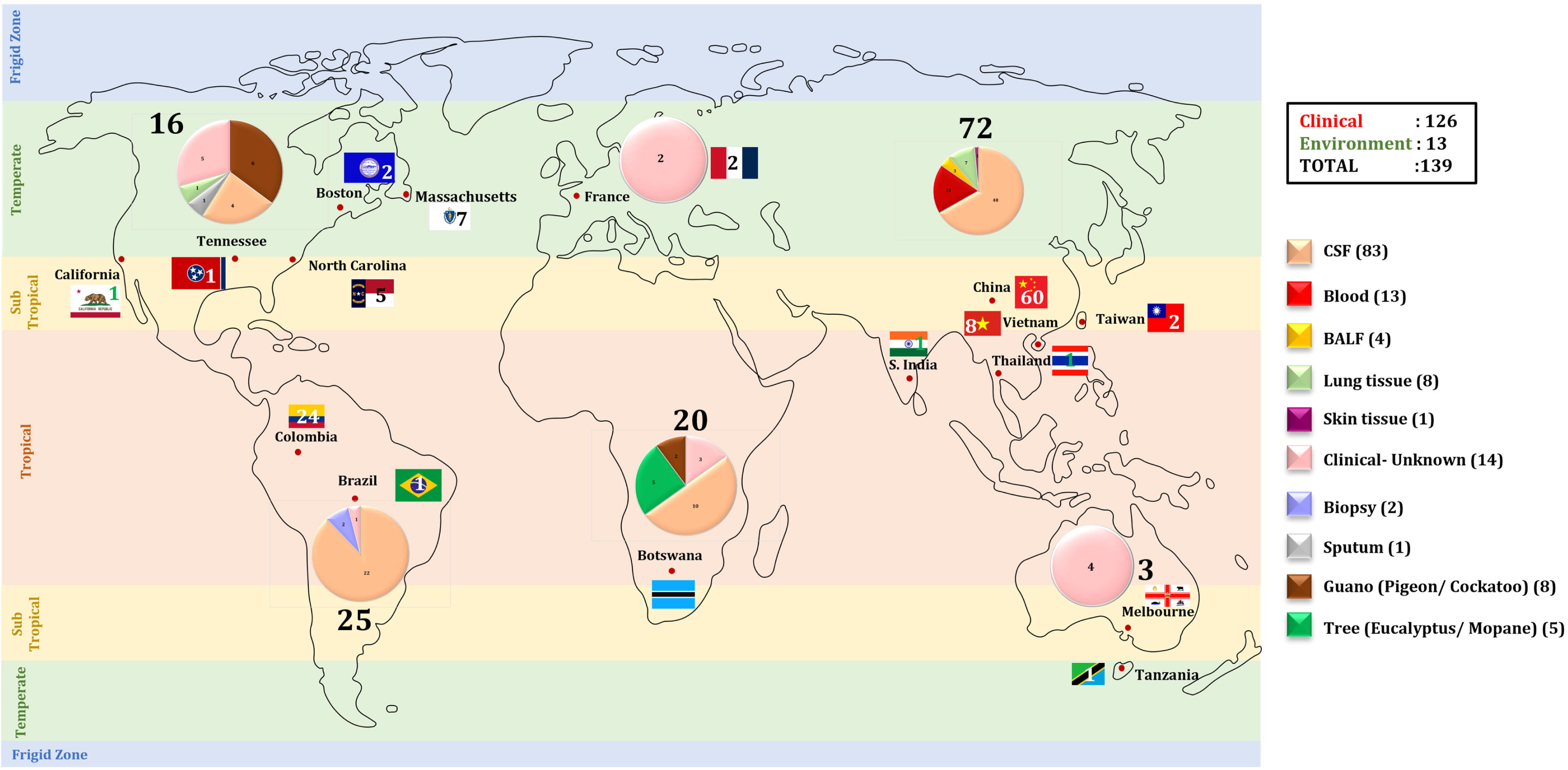
a) Geographic distribution and isolation source distribution across climatic zones: The geographic and climatic distribution map emphasizes the majority of prominent clinical isolates from temperate to tropical zones and depicts isolate numbers and source types throughout the world. Pie charts demonstrate the global propagation of pathogens in a multitude of climates by presenting sample origins, which are primarily cerebrospinal fluid (CSF) in clinical isolates, with environmental sources such as guano and tree parts displayed in lower portions.

**Figure 1:**
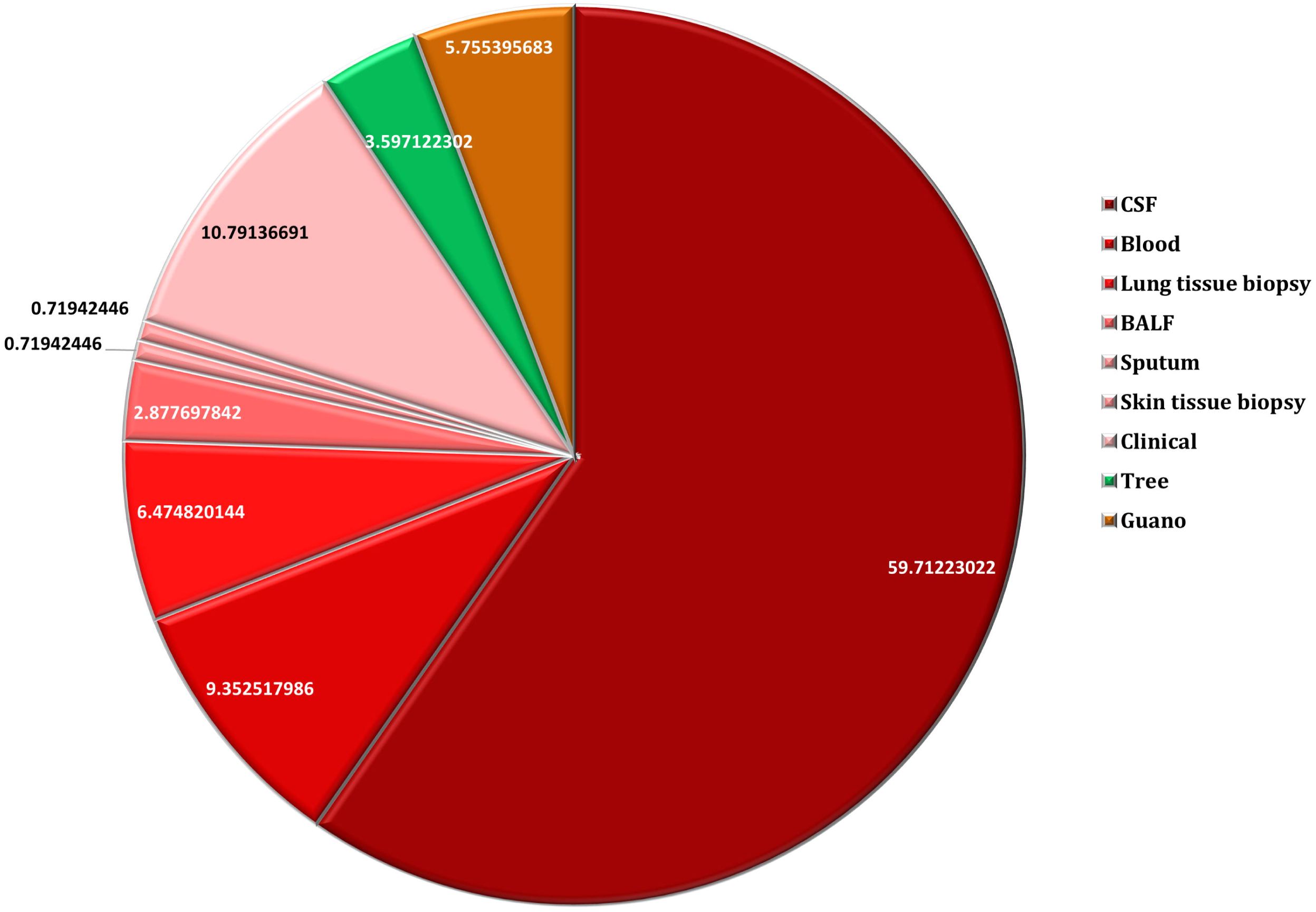
b) Distribution of isolates based on sources of isolation: The pie chart illustrates the proportional representation of C. *neoformans* isolates recovered from various sources, including clinical

### 3.2. Investigation of geographic hotspots of serotypes over the years

The majority of the genomes belonged to serotype A, which was invariably detected across all continents (N. America, S. America, Europe, S. Africa, Asia and Australia). The serotype A demonstrated sporadic distribution with the number of genomes, yet persistent distribution over nearly three decades, spanning from 1998 to 2021. This pattern indicates its long-term global prevalence and conservativeness. Whereas serotype D, the first evolved serotype, was found to be rare, represented by only a single isolate in our dataset, which was collected in 1970 (Figure 2, supplementary figure 1).

**Figure 2:**
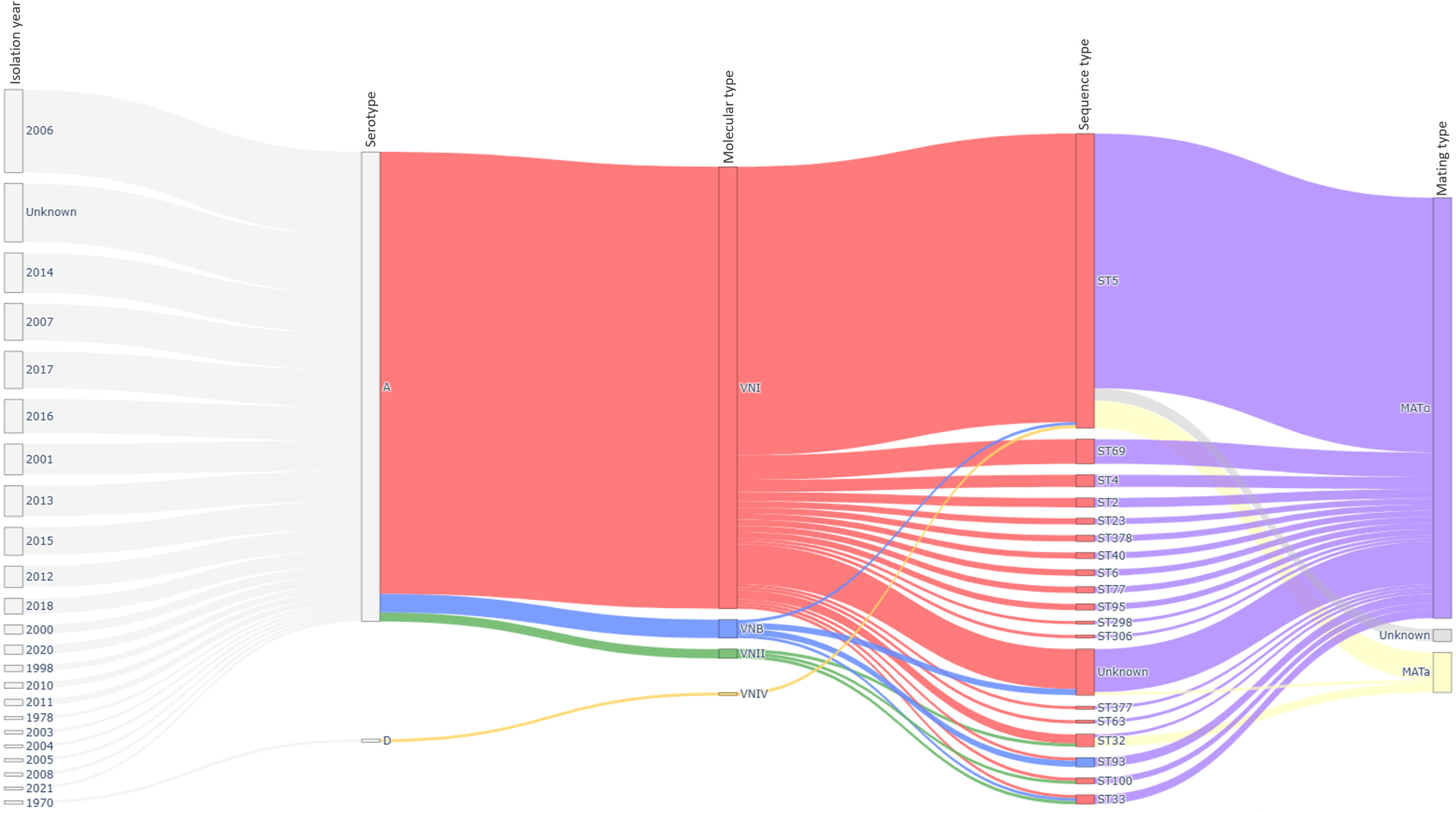
Temporal dynamics of serotypes, molecular type, ST and MAT locus: The temporal dynamics Sankey graph was plotted using Pandas for data processing and Plotly for data visualization in Python. The graph illustrates the partitioning of genomes by isolation year, serotype, molecular type and sequence type, revealing predominant VNI/ST5 MATa lineages and minor representation of other types. The flow widths indicate frequency, with large MATa dominance and rare occurrence of MATa and VNIV backgrounds in the dataset.

### 3.3. Temporal and molecular shifts in *C. neoformans* - A diverse lineage

The molecular types VNI, VNII, VNB, and VNIV were identified in *C. neoformans*, with VNI (88%) portraying the predominant lineage globally. This was followed by VNB, VNII, and VNIV. The sustained presence of VNI from 1998 to 2021 highlights its long-standing and stable circulation across time, underscoring its epidemiological dominance (Figure 2, supplementary figure 2).

### 3.4. Temporal trends and geographical spread of STs

The *C. neoformans* study genomes belonged to 19 sequence types (STs), with ST5 predominating by 63% (n = 88), accounting for 69% of clinical (CSF and blood isolates) and 54% of environmental isolates. Despite the presence of multiple lineages of STs, ST5 maintains an epidemiological niche across all reported continents, predominantly in Asia, South Africa, North America, Australia, and Europe. This data aligns with the reports that ST5 is the pandemic lineage accountable for the majority of cryptococcal meningitis cases in East and Southeast Asia. The emergence of various other STs (ST69, ST377, ST378, ST93, ST95, ST2, and ST306 appearing alongside ST5) observed from 2006 (Figure 2, supplementary figure 3).

### 3.5. Distribution of MAT genes

The present data reinforce the global distribution of MAT genes. The MATα mating type predominates across all the continents, accounting for up to 92% of total isolates. In the Asia sub-continent, MATα accounts for about 90.37%, Australia 75%, S. Africa 85%, S. America 100% and N. America 100%. Despite this skewed distribution, unisexual reproduction (MATα X MATα) is high; they still generate genetic diversity with four molecular types (VNI, VNII, VNB and VNIV). However, the predominance of VNI across the years and continents indicates that this particular type has a selective advantage over the others. The vast majority of MATα mirrors the global dominance of the α mating type, which is associated with higher virulence and predominance in clinical isolates. In some regions, like the Americas, MATa is absent, but recombination is still observed, highlighting the role of unisexual mating in maintaining diversity (Figure 2, supplementary figure 4).

### 3.6. Genome-wide diversity profiling

The genome-wide comparison heat map depicted in Figure 3 demonstrates inter- and intra-continental similarity and divergence. The intra-continental divergence of S. African isolates (n=20) exhibited the highest genomic diversity, which might be due to the high infectious rate/ host pathogen interaction that might be due to the co-evolutionary relationship between the pathogen and its host, driving genetic diversity. This confirms the fact that South Africa remains a hotspot of genetic diversity with the global *C. neoformans* population. The pathogen may adapt to evade the host’s immune system, leading to genetic changes. N. America (n=16) showed the second highest diversity, followed by S. America (n=25), Europe (n=2) and Australia (n=4). While isolates from Asia (n=72) showed minimal genetic diversity, this underscores the fact that the strains are highly clonal within the dominant lineage. The inter-continental level, more varied colour blocks in the heat map, suggesting that Asia’s population has evolved separately, with minimal recent genetic interchange with populations from other continents (Figure 3).

**Figure 3:**
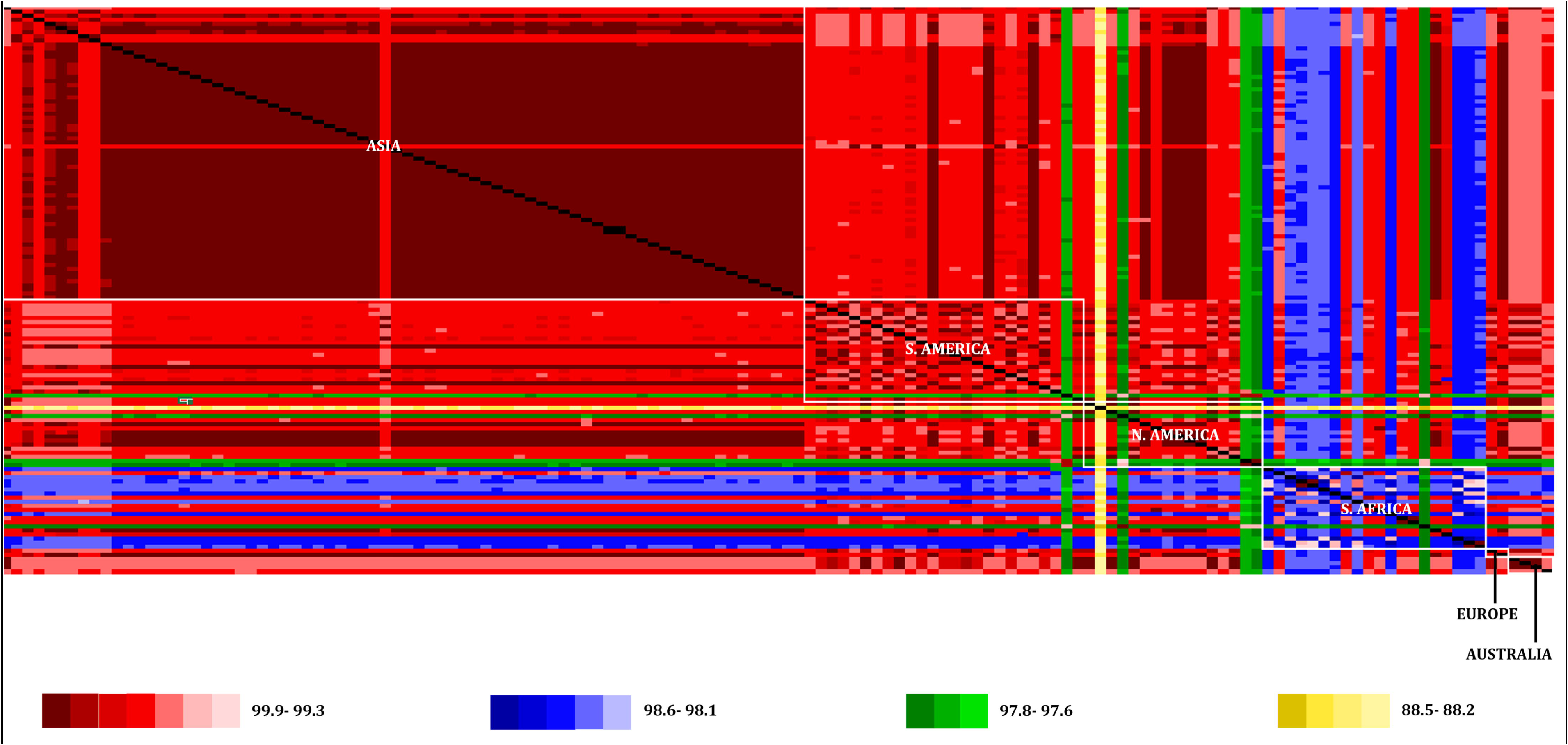
Genome-wide diversity profiling: The heatmap visualizes genome-wide nucleotide similarity among isolates from different continents, revealing distinct regional clusters with color-coding for genetic similarity and divergence. Diagonal black blocks show 100% similarity, while red (99.9-99.3%), blue (98.6-98.1), green (97.8-97.6%), and yellow (88.5-88.2%) bands indicate zones of divergence, or outlier lineages in the global dataset.

### 3.7. Whole genome SNP profiling

The dendrogram in Figure 4 reveals clear substructure, representing the pairwise SNP distances between the isolates. The isolates cluster into several major groups, with a dominant large cluster and a few genetically distinct outliers. The percentage of SNPs fluctuates from 0.0019% to 0.014%, indicating relatively low genetic diversity among the isolates, which is in concordance with genome-wide diversity profiling.

**Figure 4:**
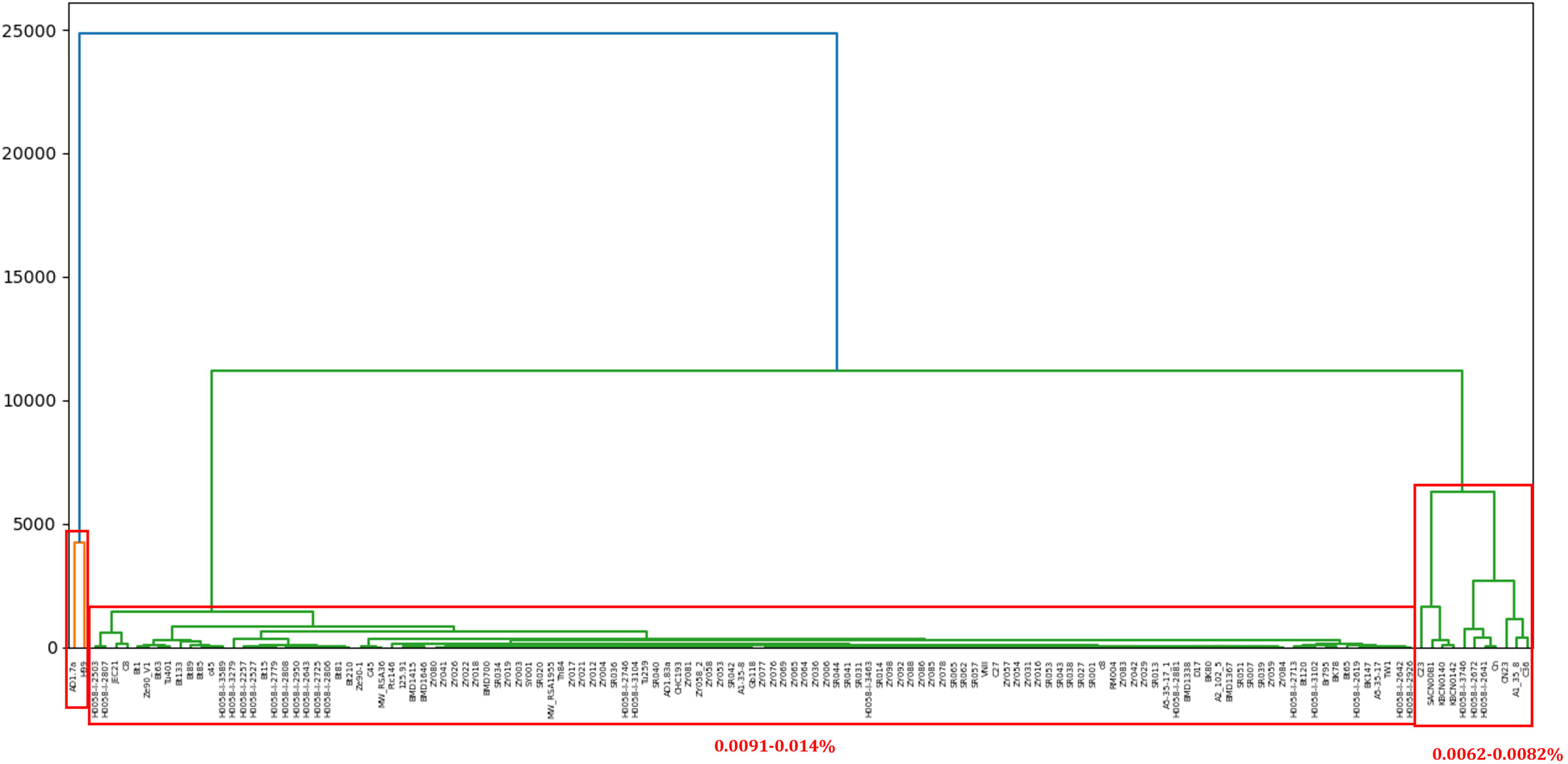
Whole-genome SNP profile: The whole-genome SNP-based phylogenetic dendrogram illustrates the genetic similarity and evolutionary divergence among the isolates.

### 3.8. Lineage analysis and population stratification

The phylogenetic analysis was made based on the MLST and orthologous genes. In both the trees, the midpoint separated the isolates into two major clades: *C. neoformans* var. *grubii* (clade 2) and *C. neoformans* var. *neoformans* (clade 1), corresponding to the serotypes A and D. Clade 2, belonging to serotype A, was subdivided into various subclades based on molecular types. Unlike the phylogenetic tree constructed with MLST genes, the tree constructed with orthologous genes showed a clearer and more distinct delineation of molecular types. VNI is a major global clade of serotype A, while VNII is the smaller and less diverse subclade of serotype A. VNB is the highly diverse subgroup of serotype A, which is distinctly separated from VNI and VNII. The orthologous gene-based phylogenetic trees were depicted in Figure 5. The present investigation emphasises that for high resolution of phylogenetic analysis, genome-scale markers should be used rather than classical MLST.

**Figure 5:**
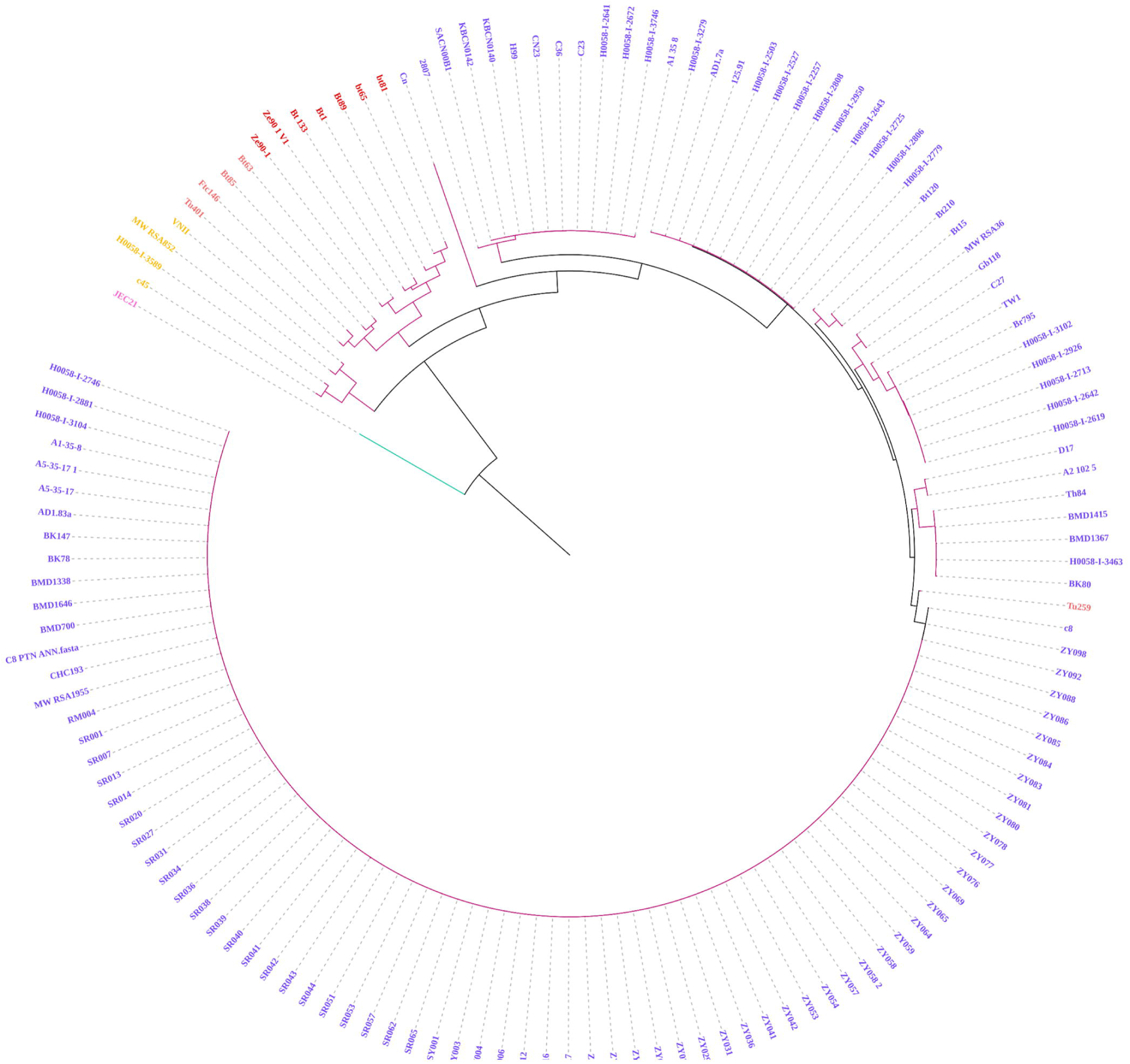
Orthologous gene-based phylogenetic tree: Orthologous gene-based phylogeny was constructed by the maximum-likelihood method with MegaX. 1000 bootstrap replicates were calculated to assess robustness. The phylogenetic tree was divided into two major clades, six subclades, and further delineated into many clusters of C. *neoformans.* The serotype D is in clade 1, and the serotype A is in clade 2. Clade 2 was further divided into subclades and clusters were based on molecular types. The phylogenetic data were visualized using iTOL

### 3.9. Pan-Genome analysis

Pan-genome analysis facilitates the understanding of the genome architecture (Figure 6a). The *C. neoformans* pan-genome was constructed with 6665 orthogroups containing 841606 genes. The 6665 orthogroups were subdivided into core (62.5%; 3780 orthogroups with 526244 genes), shell (37.4%; 2856 orthogroups with 315261 genes) and cloud orthogroups (0.01%; 29 orthogroups with 101 genes). The core orthogroup encompasses a single-copy orthogroup (n=2902) and a multiple-copy orthogroup (n=878). In addition, 378 unassigned genes (new genes) contributing to 0.04% of the total genome were identified as novel genes that did not cluster into any orthogroup, reflecting unique lineage-specific innovations (Figure 6b). Integrating the functional annotation of these orthogroups with virulome and resistome datasets provided functional insights into genomic specialisation. Notably, eleven virulence genes belong to the core genome and seven belong to the shell genome, suggesting that most virulence determinants are highly conserved but exhibit some variability. In contrast, resistome analysis revealed that all the genes involved in the resistome profiling belong to the core genome, which aligns with the concept that cryptococcal resistance arises via SNPs rather than gene gain/ loss. Together, these findings highlight that while the *C. neoformans* genome exhibits flexibility through accessory gene content, with virulence traits being partly conserved and partly variable, resistance mechanisms are strictly conserved within the core genome. This distinction underscores how virulence and resistance genes are subject to different selective pressures: Virulence genes are largely conserved to maintain pathogenic fitness, whereas resistance genes evolve in response to environmental or drug-induced stress.

**Figure 6:**
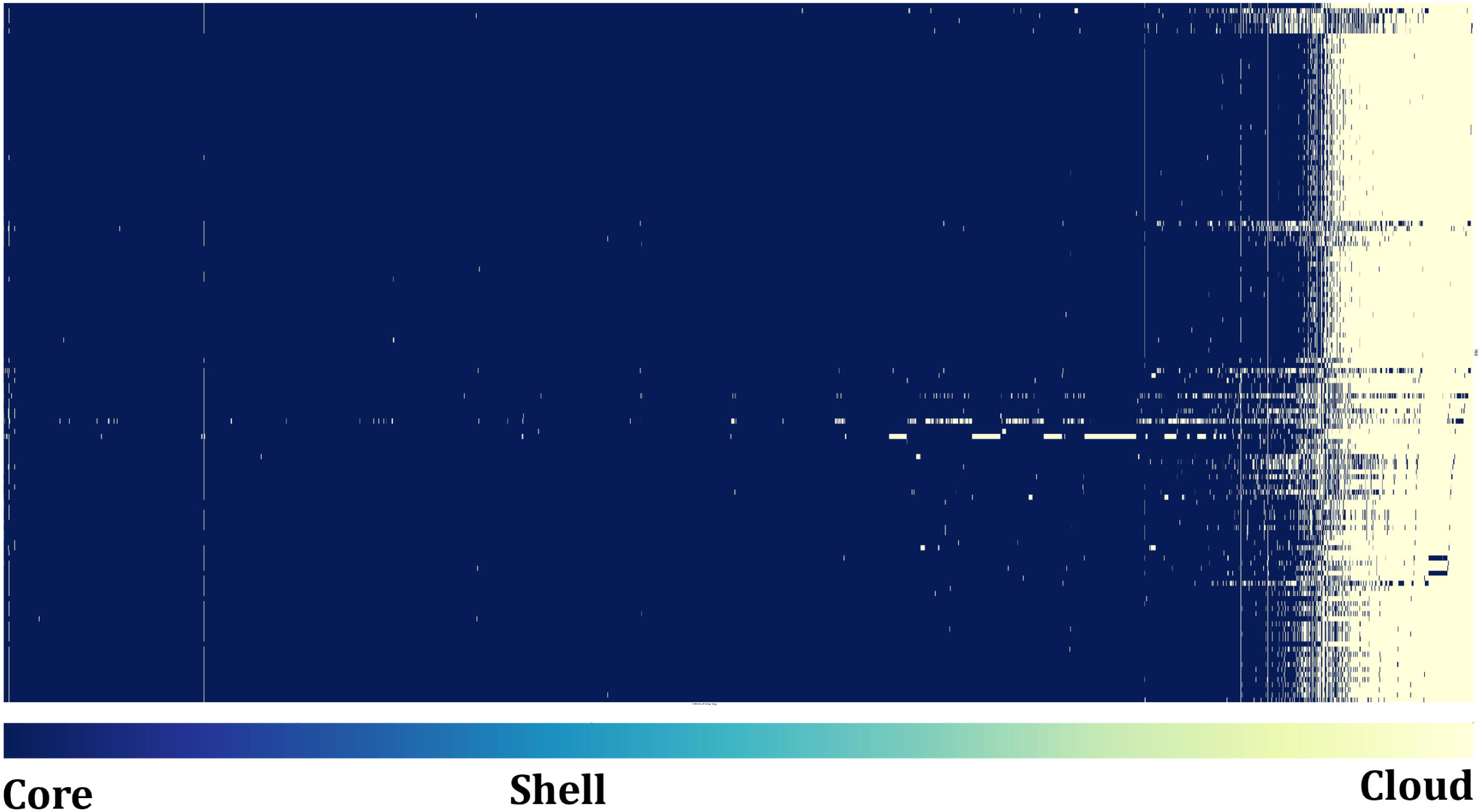
Pangenome analysis: a) Heat map of Pangenome analysis: The distribution of core, shell and cloud genes among 139 C. *neoformans* strains. The heat map was generated using Matplotlib, Pandas and Seaborn in Python.

**Figure 6.**
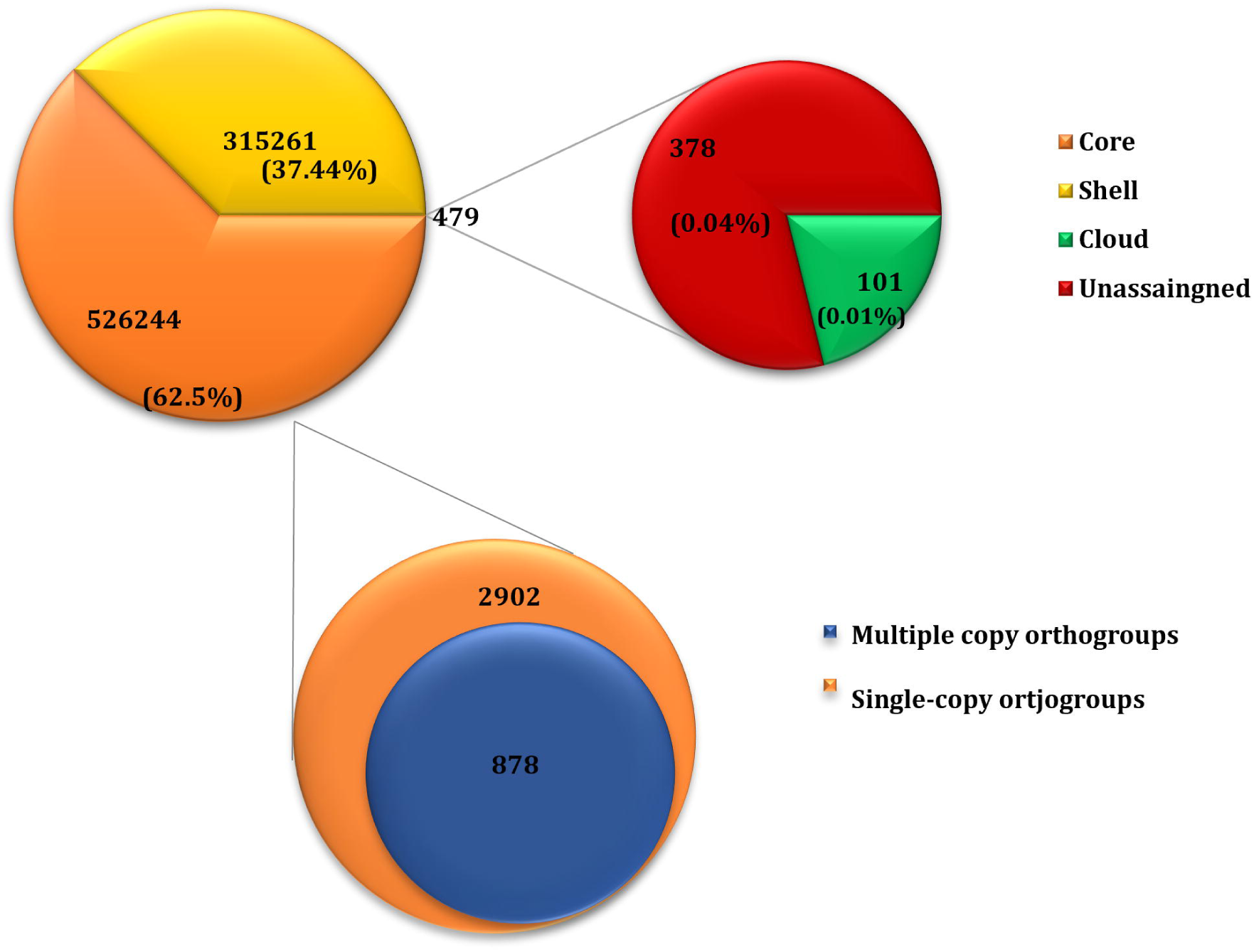
b) Pie chart of Pangenome analysis: Pie chart presenting the Core (single and multi-copy core orthogroups), Shell, cloud orthogroups

**Figure 6.**
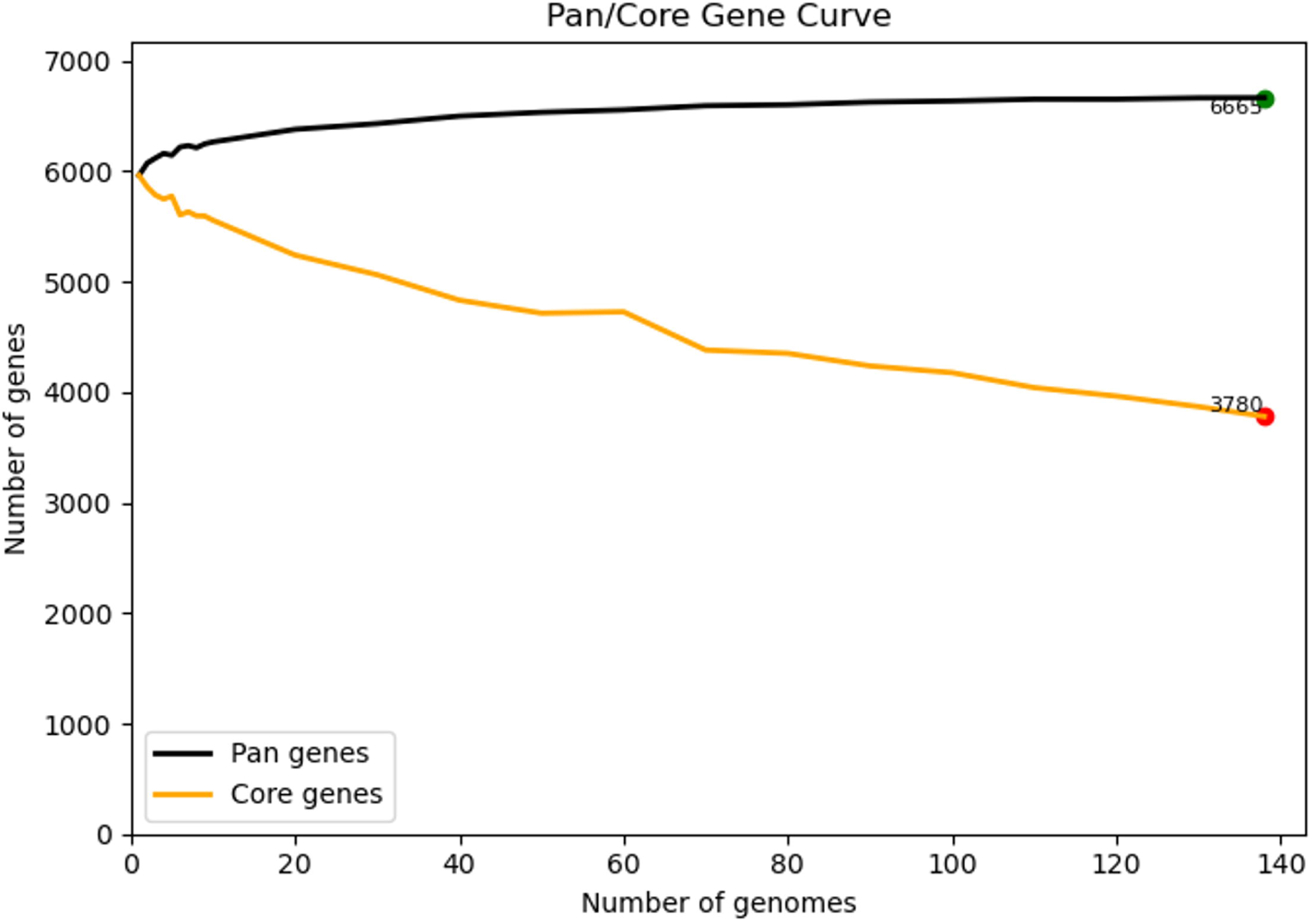
c) Closed pangenome structure: The pangenome curve demonstrates the closed pangenome nature emphasized by the pangenome curve saturated with the addition of new genomes, that is, consistent with limited acquisition of new genes among isolates.

### 3.10. Comparative Virulome Profiling

We have identified 274 virulence genes, out of which 17 important virulence genes were selected based on their degree of pathogenicity^27^ (Figure 7). Out of 126 clinical strains, only 17 strains have all the VRGs. 80% of the CSF isolates are VRG-positive strains, suggesting that these strains have high pathogenic potential to disseminate and survive in the central nervous system. These highly virulent strains belong to ST5 (53%), and Serotype A. Both the clinical and environmental isolates were found to have genes mostly associated with capsule synthesis (*CAP10*, *CAP59*, *CAP60* and *CAP64*), melanin synthesis (*LAC1* and *LAC2*), iron acquisition genes (*CFT1*, *SIT1*, *CIR1* and *RIM101*), mannosyl transferase genes (*ALG3* and *MPD1*), superoxide dismutase genes (*SOD1* and *SOD2*), Urease gene (*URE1*), phospholipase gene (*PLB1*) and protease gene (*PIM1*), highlighting their conserved role in virulence. The pattern of gene loss appears scattered, rather than confined to a single phylogenetic clade, indicating that the gene loss isn’t strictly lineage-specific, but rather sporadic across the population

**Figure 7:**
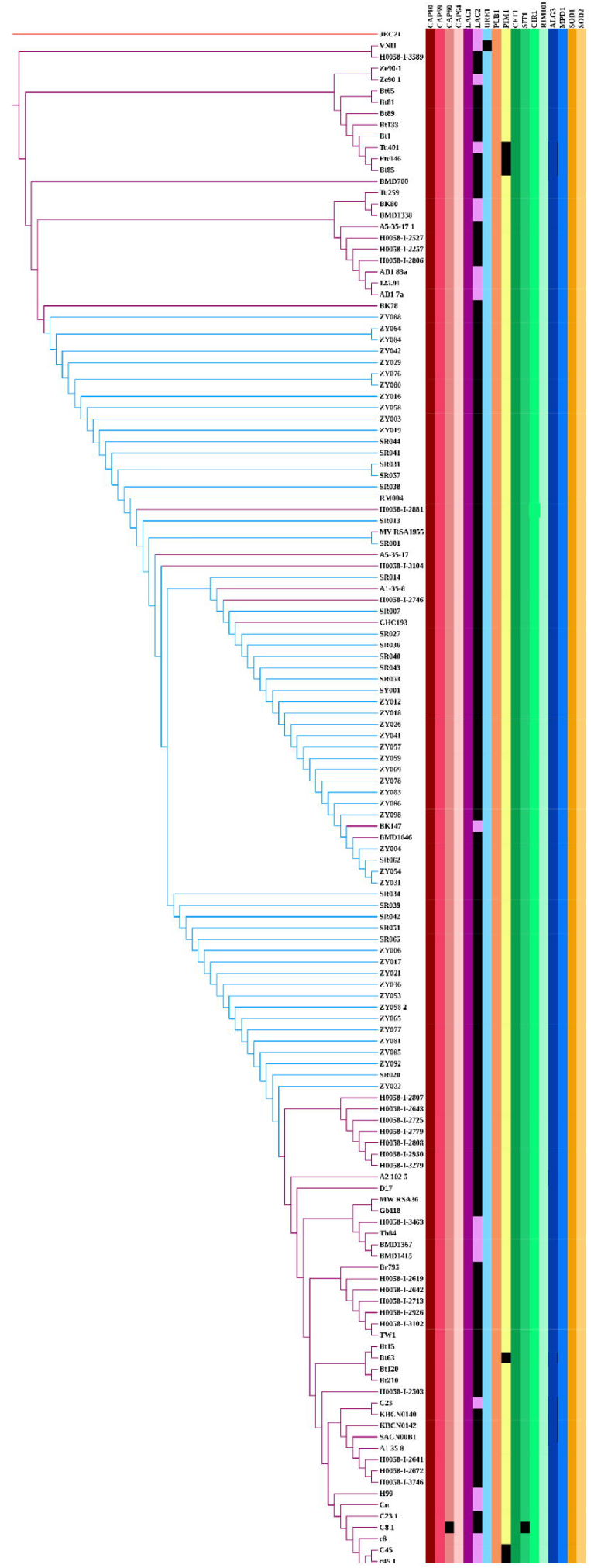
Virulome of C. *neoformans:* The major virulence genes (n=l 7) in C. *neoformans* are screened among the 139 study genomes. The coloured regions represent the presence of genes and the black colour represents the absence of genes. The data was visualized using iTOL.

### 3.11. Comparative Resistome Profiling

In *Cryptococcus*, the development of resistance is typically driven by point mutations in the conserved essential genes, aneuploidy and mere presence or absence of genes^28^. The major genes studied for mutations include *ERG11* (Lanosterol 14α demethylase), *TUB1* (βtubulin), *FKS1* (β-glucan synthase), *UXS1* (UDP-glucuronate decarboxylase 1), *FUR1* (Uracil phosphoribosyl transferase), and *FCY2* (Cytosine permease), which are associated with azole resistance, echinocandin resistance, and pyrimidine analogue resistance, respectively. The presence (overexpression) or absence of *AFR1, NRG1* and *MSH2* also plays a vital role in promoting antifungal resistance^28,29^. The major mutations studied in *ERG11* include Y145F, I99V, G484S, and G470R. The mutational sites studied in *TUB1* include H6Q and F200. The mutational sites studied in *UXS1* include D306C and Y217C. The mutational site studied in *FKS1* includes S643P. The mutational sites studied in *FUR1* include P140S, G87D and G190E. The mutational sites studied in *FCY2* include A157V. The analysis shows that all of the study isolates (n=139) depicted mutations in *ERG11* and *FUR1,* highlighting the resistance to azoles and pyrimidine analogues and sensitivity to polyenes. The same result has been proved in the *in vitro* antifungal susceptibility study and MIC conducted using the azoles and the available study isolates H99 and Cn. The comprehensive Figure 8a, 8b, 8c and 8d and Table 1 illuminate an overview of antifungal resistance in *Cryptococcus neoformans*, integrating phylogenetic context, resistance gene profiling and *in vitro* drug susceptibility, thereby highlighting the clinical relevance of drug resistance mutation for effective therapeutic options.

**Figure 8:**
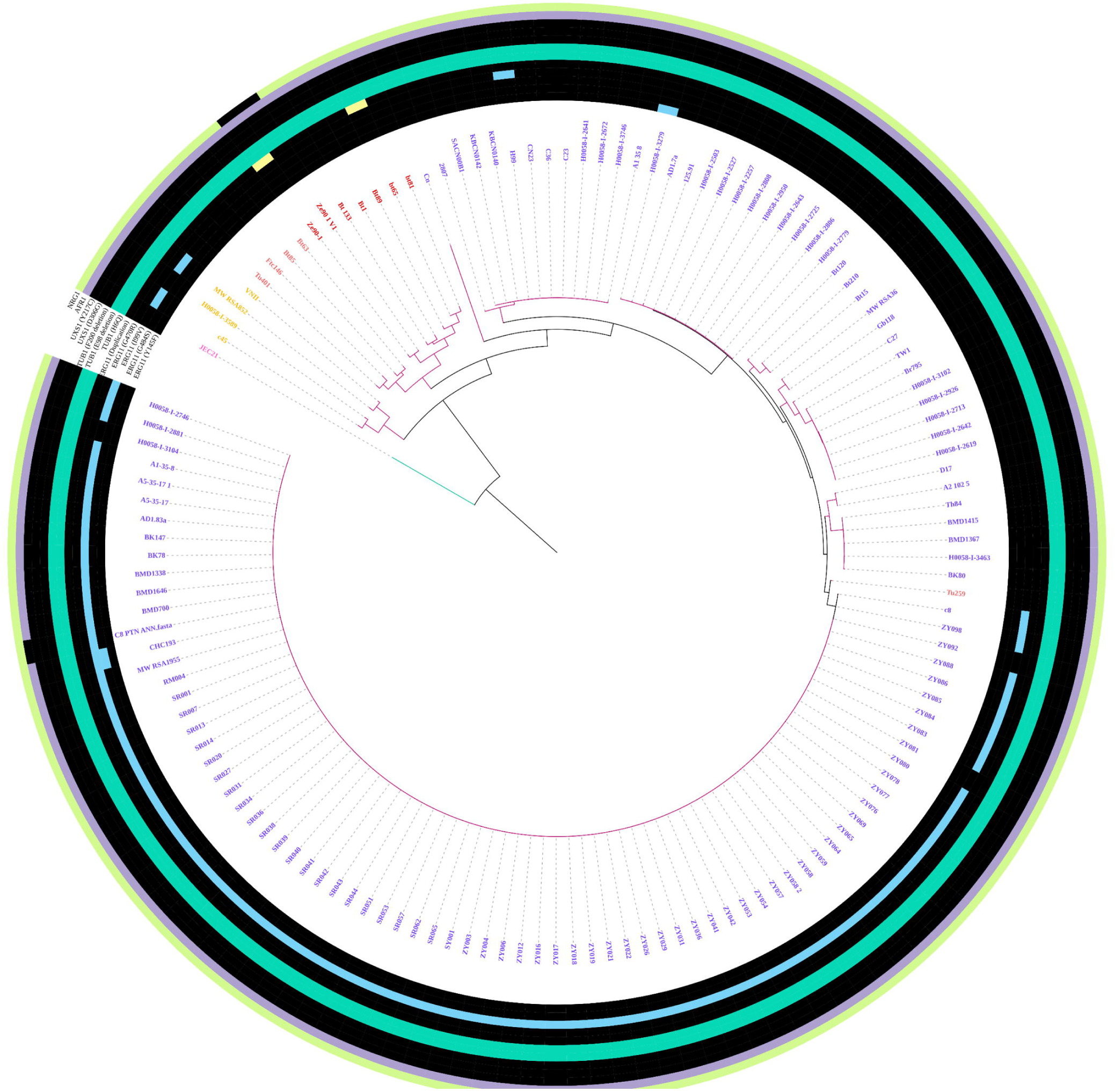
Resistome of C. *neoformans:* a) Azole resistance: Mutations in *ERGll* (n=4), Aneuploidy of *ERGll* (n=l), *AFR]* (n=l), indirect mutations in *TUB]* (n=3), *UXSJ* (n=2) and *NRG1* (n=1). The coloured regions represent the presence of genes and the black colour represents the absence of genes absence of genes

**Figure 8:**
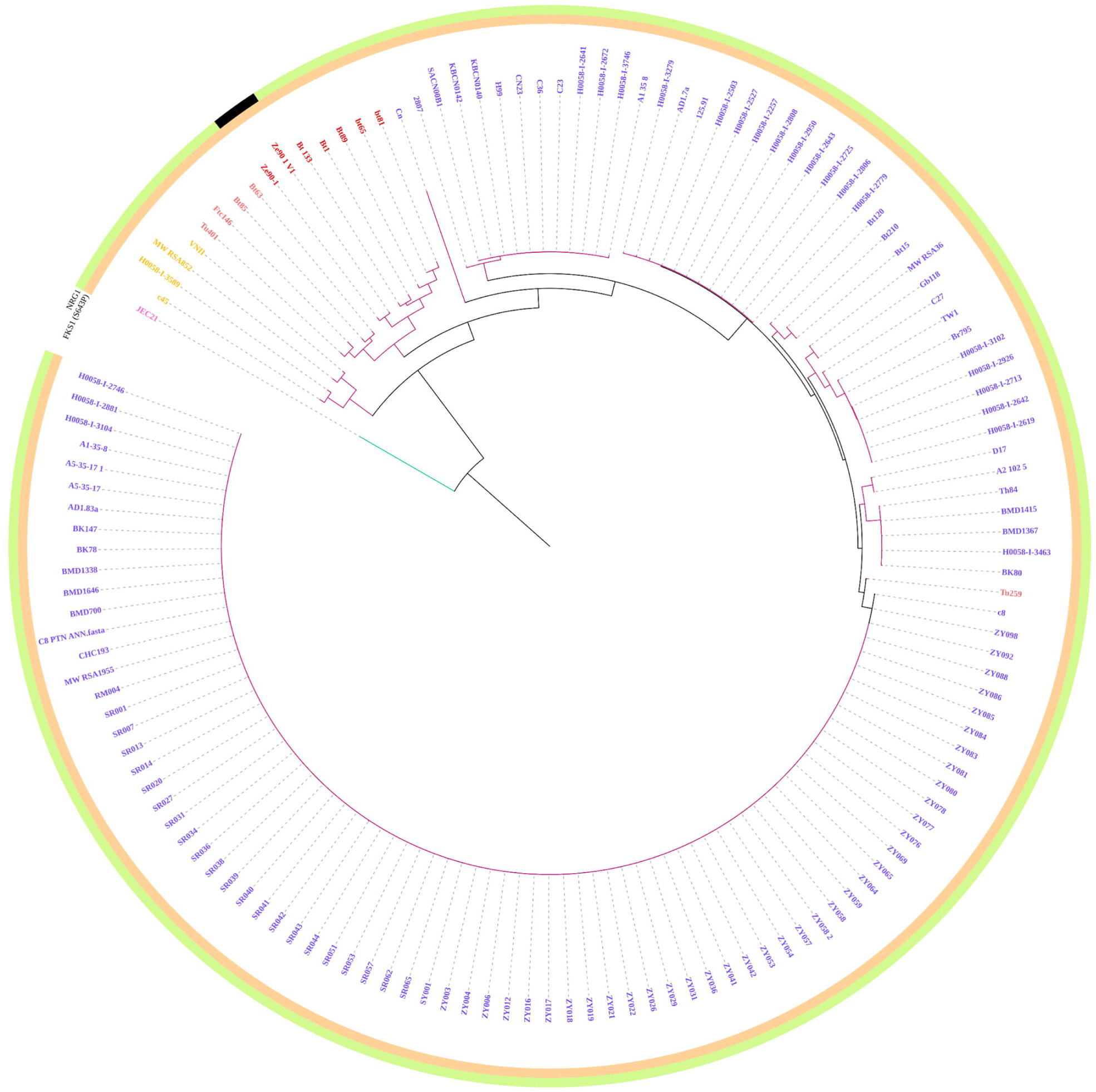
Resistome of C. neoformans: b) Echinocandin resistance: Mutations m FKSl (n=l) and NRG] (n=l). The coloured regions represent the presence of genes and the black colour represents the absence of genes

**Figure 8:**
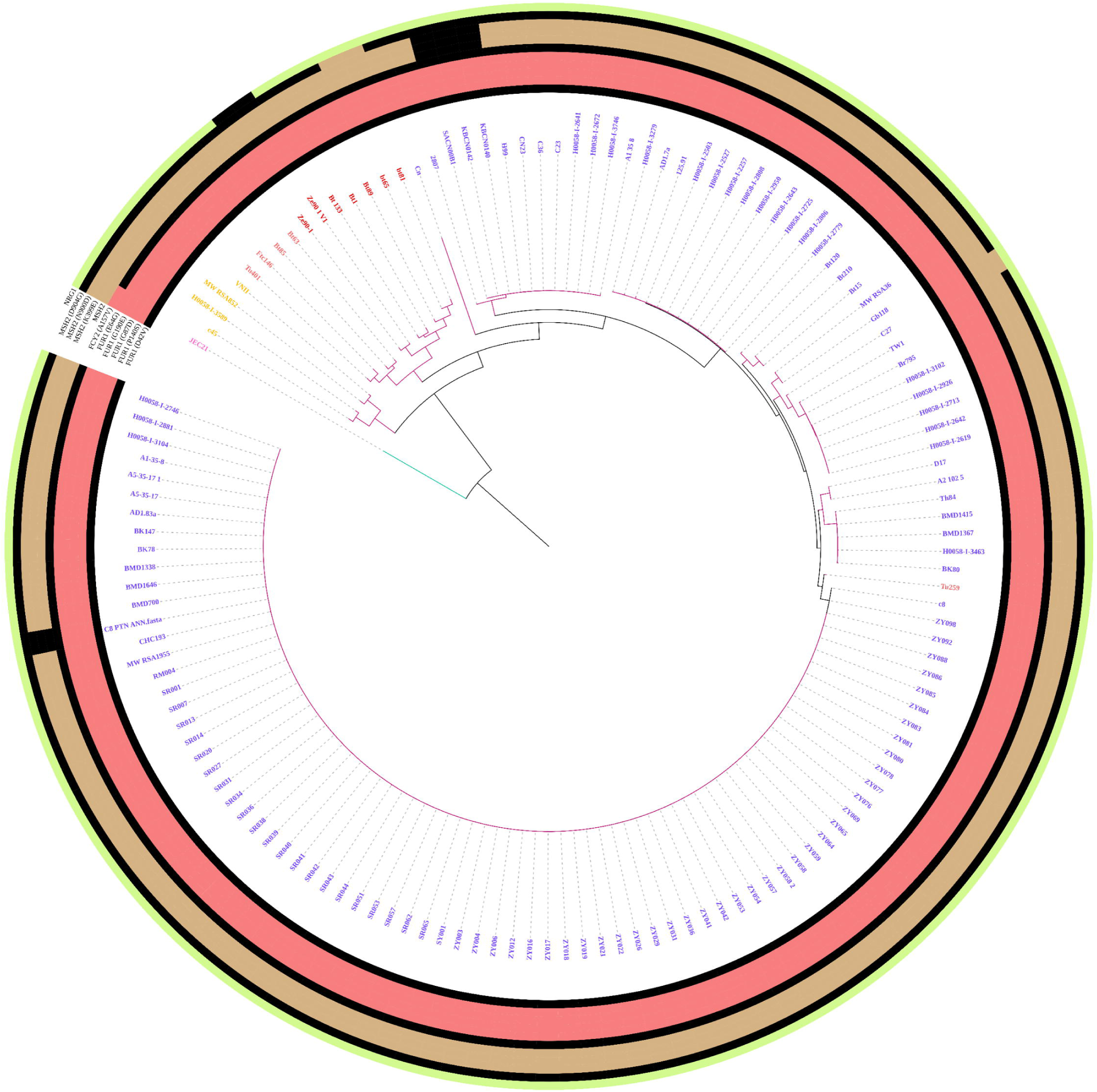
Resistome of C. *neoformans:* c) Pyrimidine analogue resistance: *FUR]* (n=5), *FCY2* (n=l), *MSH2* (n=5) and *NRG]* (n=l). The coloured regions represent the presence of genes and the black colour represents the absence of genes

**Figure 8.**
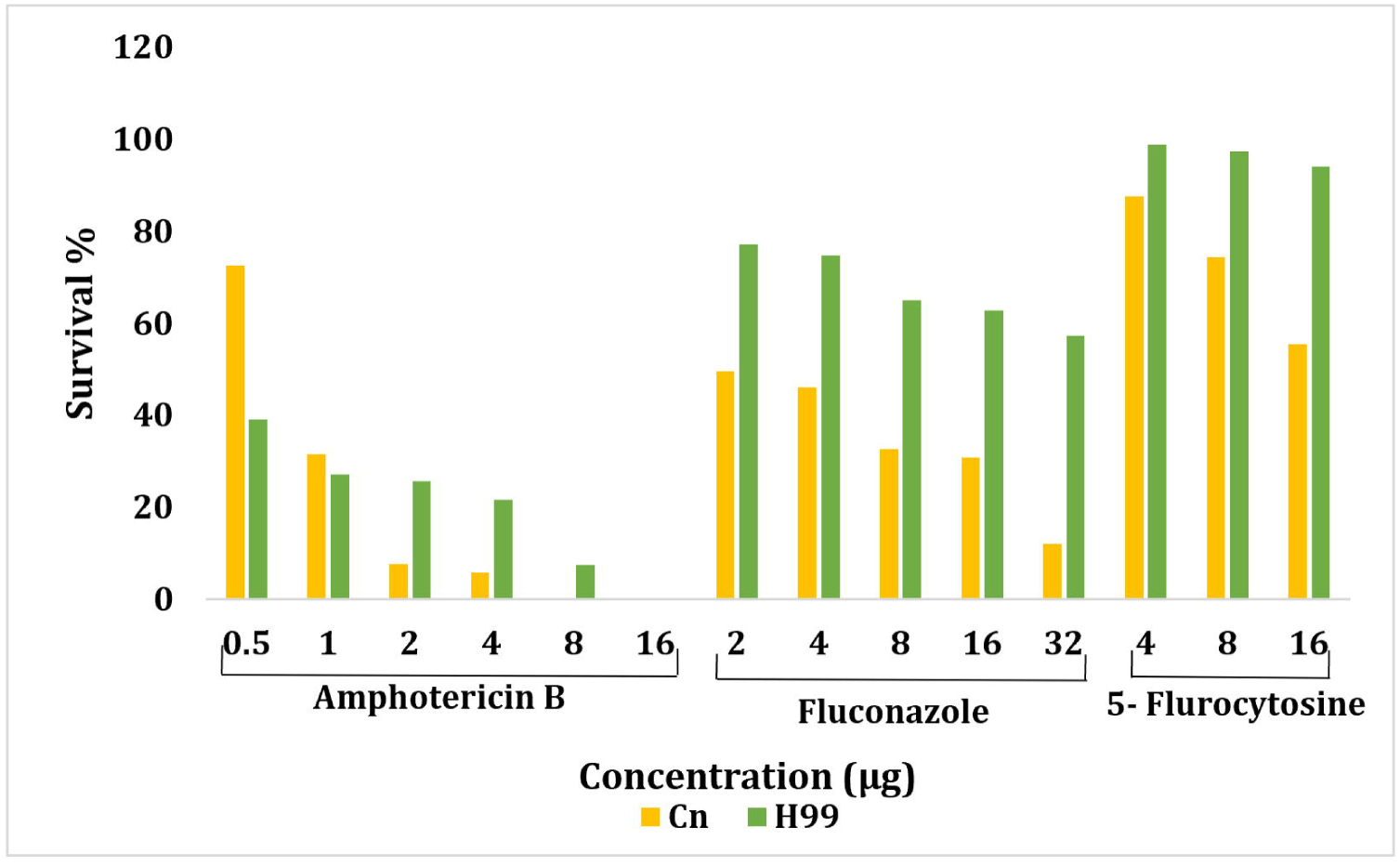
d) Minimum Inhibitory Concentration assay: Fluconazole, Amphotericin B and 5-fluorocytosine tested against Cn and H99 study strains. The data was visualized using iTOL.

**Table 1:** AFST of Voriconazole, Itraconazole and Ketoconazole against Cn and H99 strains

| S.no. | Antifungal Drug | Conc. (µg) | ZOI (mm)<br>(*C. neoformans* H99) | R/I/S | ZOI (mm)<br>(*C. neoformans* Cn) | R/I/S |
| --- | --- | --- | --- | --- | --- | --- |
| 1. | Voriconazole | 10 | 26 | S | 50 | S |
| 2. | Itraconazole | 10 | 11 | I | 12 | I |
| 3. | Ketoconazole | 10 | 23 | R | 26 | R |

S= Susceptibility
I= Intermediate
R= resistance

## 4. Discussion

This comprehensive comparative study of *C. neoformans* genomes collected during 1970-2021, analysing 139 publicly available strains from diverse environments and host factors worldwide. The study examined the origin of strain, year of isolation, ST, serotypes, molecular types, mating type, resistome, virulome and pangenome analysis. Environmental reservoirs, including pigeon guano, cockatoo excrement, mopane and eucalyptus trees, reinforce the zoonotic-ecological transmission cycle. The year-wise distribution of the isolates aligns with the epidemiological data, indicating a rise in cases during the mid-2000s, from which 41% isolates were collected. Notably, nearly half of these collected strains were from Botswana, a recognised diversity hotspot within Sub-Saharan Africa^30,31^.

The temporal dynamics study reveals key findings on the molecular epidemiology of *C. neoformans*. ST5 is a dominant lineage, widespread among the VNI molecular type^32,33^. ST5 is globally dominant in cryptococcal meningitis, suggesting clonal expansion and international dispersal linked to migration, urbanisation, and ecological flexibility. Different STs exhibit distinct geographic distributions, with ST69 (the second predominant ST, VNI, Serotype A) and ST2 being common in South America, particularly in Colombia. ST5 is primarily associated with the VNI molecular type and serotype A. Notably, South America shows high diversity with 12 different STs. The MATα locus is predominant globally^34^. Its predominance is notably found among VNI and VNII molecular types and serotype A, while MATa is less common, found among VNB molecular type serotype A. MATa is notably present in Botswana (S. Africa) and China (Asia), which could be of epidemiological and evolutionary interest as they suggest opportunities for sexual reproduction and potential genetic recombination. Clinical isolates closely mirror those from environmental niches like pigeon guano, reinforcing the zoonotic-ecological cycle of *C. neoformans*^35^ (ST5, VNI isolates from both clinical and avian sources in N. America, S. Africa and Asia) (Figure 9).

**Figure 9:**
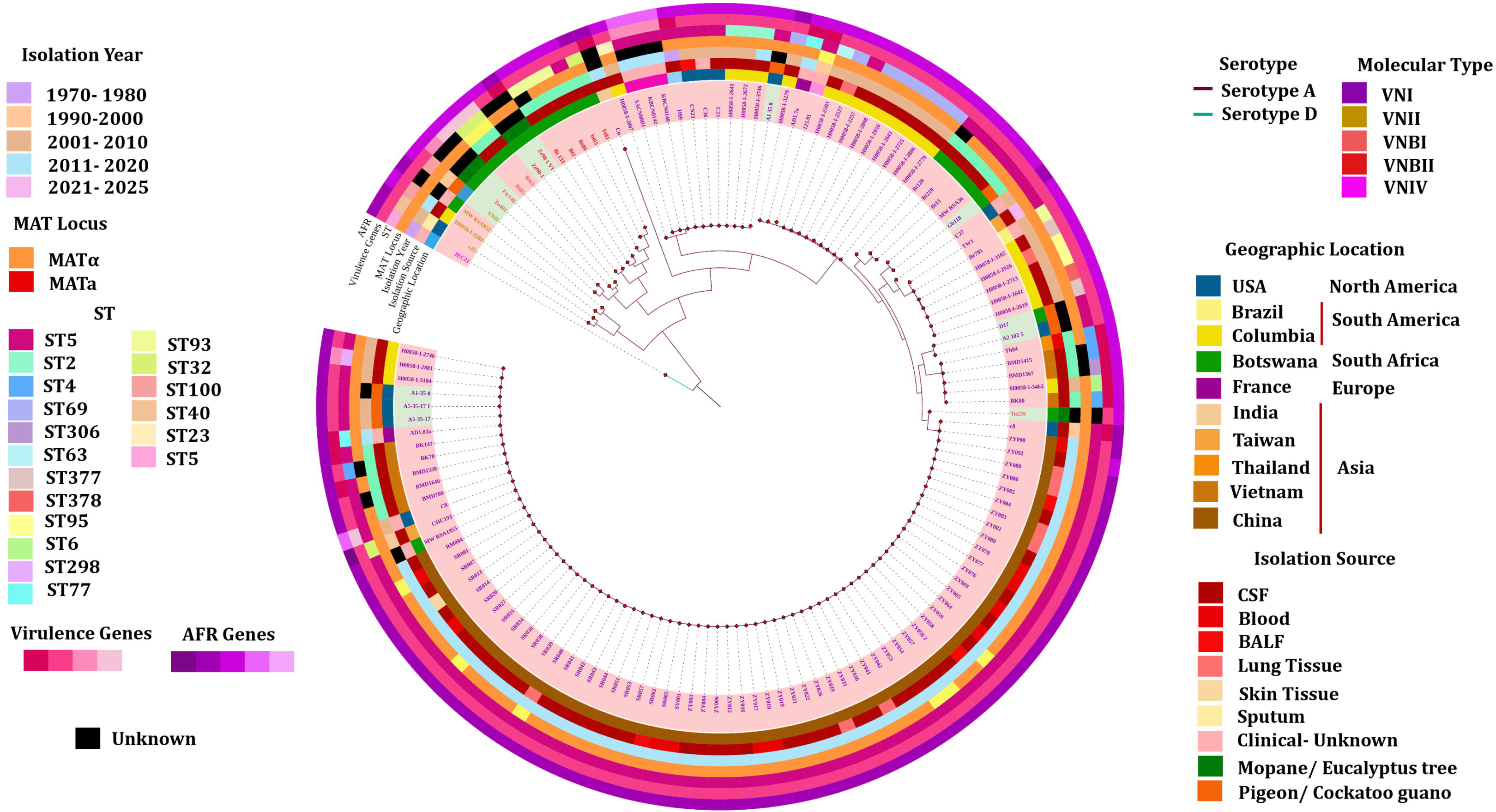
A complete depiction of Orthologous gene-based phylogeny: The tree consists of geographic location, sources of isolation, temporal dynamics of serotypes, molecular types, STs, mating type, number of virulence genes and number of mutations in AFR genes, that were prevalent among the study genomes. The data was visualized using iTOL.

The genetic divergence analysis reveals a dominant clonal population of *C. neoformans* var. *grubii* across Asia, S. America and Europe (intracontinental analysis) with an explicit genomic shift between Asian and non-Asian isolates (intercontinental analysis). Also, it was noted that the molecular type VNB is present only in S. African genomes and a study conducted in 2017 on tracing the genetic exchange of biogeography of *C. neoformans* stated that VNB is predominantly found in S. Africa^36^. Studies have reported the presence of VNB isolates in S. America, stating a possible transmission of VNB isolates from S. Africa to S. America^10^. The high nucleotide similarity and limited number of SNPs suggest that this pathogen may have originated from a recent clonal expansion, with unisexual mating contributing to the limited evolutionary events^37^. The ability of *C. neoformans* to thrive in diverse environments and host conditions may also play a role in its widespread distribution^36^. While the genetic divergence analysis emphasises genetic variation across the population, the orthologous gene-based phylogenetic tree construction underlines ancestral branching patterns among lineages. This highlights the deep divergence of VNII, VNB and VNI, suggesting that they may represent an ancestral or highly diverse lineage within serotype A. The clustering shows clear evolutionary separation between serotypes A and D, supporting the well-established molecular phylogeny of *Cryptococcus neoformans* var. *grubii.* Serotype D forms its own isolated cluster, distinct from all A-lineages.

Pan-genome analysis indicates that *C. neoformans* has a closed pangenome over time (Figure 6c). Though the total number of orthogroups continues to increase with the addition of new genomes, the rate of new gene acquisition is lower, indicating genomic stability within the *C. neoformans* var. *grubii*. Notably, most of the virulence (65%) and resistance (88%) genes belong to the core genome, suggesting their conservation. The 37% of accessory genomes highlights the genetic flexibility that enables environmental adaptation and occasional emergence of antifungal resistance.

The conservation of primary virulence genes (*CAP* genes, *LAC1*, *SOD* genes, *PLB1*) in the core genome indicates the essential role of retaining these traits, likely due to their essential roles in establishing infection within the host. However, the virulence genes harboured in the accessory genome *LAC2*, *SIT1*, *URE1*, *PIM1*, *SOD1* and *MPD1* were observed to have sporadic gene loss events, suggesting local environmental adaptations or loss of selective pressure in specific ecological niches. This variation in gene conservation may contribute to differences in infection severity and epidemiological patterns. The sporadic, occasional absence of genes, often not clustered within a single branch but scattered, reveals independent gene loss events across phylogeny. Infrequent gene loss events do not appear restricted to one lineage or clade, indicating convergent or parallel loss rather than an ancestral event lost by an entire lineage. Such losses may reflect local environmental adaptation, mutation or perhaps loss of selective pressure in specific ecological niches. The high conservation of virulence genes signifies a strong evolutionary benefit. The occasional loss of certain accessory virulence genes could result in variable virulence among strains, however not lose its infection severity

In *C. neoformans*, azole resistance arises through a multifaceted defence strategy that involves *ERG11* point mutations and *ERG11* aneuploidy, increased expression of efflux pumps such as *AFR1*, and cross-resistance driven by point mutations in *TUB1* and loss of *NRG1.* The point mutations in *ERG11* result in the development of moderate resistance, whereas the mutation in *MSH2* and *ERG11* duplications result in the development of rapid resistance^38^. The point mutations in *ERG11* are I99V, G484S, Y145F and G470R^39^. The point mutation I99V is not present in the binding pocket but lies in the α-helix near substrate entry or structural elements that stabilise the active site. It was a conservative hydrophobic change that resulted in the subtle reorientation of neighbouring helices, which causes an indirect change in the geometry of the binding pocket. I99 is close to the drug access channels in CYP51. This mutation narrows or alters the dynamics of this channel (Conformational shift), limiting azole entry. But this confers low-level resistance by weakening azole binding, when combined with other mutations, rising the MIC significantly. This mutation confers resistance-associated functional alteration. The point mutation G484S was present near the heme and azole binding cavity. This mutation disrupts the local packing and introduces polar side chains near the azole binding cavity altering the heme co-ordination reducing the drug efficacy. This mutation confers resistance associated with functional alteration. The point mutation Y145F is a major mutation that takes place close to the active site binding pocket. The tyrosine is an aromatic amino acid with a polar -OH group, whereas phenylalanine is also an aromatic amino acid, yet it lacks the polar -OH group. This eliminates the potential H-H-bonding interaction within the binding pocket, altering the local polarity significantly. This mutation is strongly associated with fluconazole resistance, as the Y145 normally coordinates with the triazole ring of fluconazole via interactions with the heme group. The Y145F makes the binding pocket more hydrophobic by significantly reducing the fluconazole affinity. This mutation also confers resistance associated with functional alteration. The mutation G470R is located in the heme-binding domain. The G470 being replaced with a bulky, charged side chain distorts heme positioning by disrupting the local helix dynamics, altering the geometry of the binding channel. The introduction of such bulkier groups narrows or blocks the part of the channel, making it harder for azoles to access the heme pocket efficiently. The electrostatic interferences of positively charged groups repel azoles or disrupt favourable polar interactions. This resistance is associated with functional alteration. Interestingly, I99V was widely present in VNI in about 53% of isolates and VNII in about 60% of isolates. It was absent in the VNIV and VNB isolates. The point mutation G484S, Y145F and G470R shows scattered presence in VNI isolates alone. The *ERG11* aneuploidy causes the overproduction of the target enzyme, reducing the drug’s efficacy. *ERG11* aneuploidy is classically described as a reversible adaptive mechanism under fluconazole stress. It is dynamic and reversible under antifungal treatment. Among all the study strains, *ERG11* aneuploidy was observed in VNB isolates. The *ERG11* aneuploidy observed in environmental strain may be suggestive of the fact that this genomic state may persist beyond transient drug exposure or selective pressures (clinical antifungal therapy and environmental azole contamination) or compensatory adaptations that reduce the fitness cost of aneuploidy, underscoring its potential role in the natural evolution of antifungal resistance in *C. neoformans*.

The mutations in the *TUB1* gene (α-Tubulin) indirectly promote azole resistance by functional alteration of microtubule modulation. The predominant mutations H6Q and E98 deletion were present in all the study isolates, whereas the mutation F200 deletion was absent in all the study isolates. The H6 region of the *TUB1* protein contributes to protein folding, stability and dimerisation with the β-tubulin, essential for microtubule assembly. The H6Q was a missense substitution that slightly alters the microtubule dynamics and its interaction with microtubule-associated proteins (MAPs). Histidine is amphipathic, which can switch between protonated and neutral states at physiological pH, useful for dynamic structural interactions. Glutamine is polar, which lacks the same protonation versatility as histidine, resulting in reduced flexibility in local interactions. These mutations are associated with functional alteration of resistance. The *TUB1*, E98 deletion refers to the loss of glutamic acid at position 98, close to the GTP-binding interface and the region that interacts with β-tubulin, critical for microtubule polymerisation and stability. The loss of glutamic acid at the 98^th^ position alters the conformation and charge distribution, potentially destabilising microtubule assembly, resulting in loss of sensitivity to azole-induced stress by shifting cytoskeletal dynamics. Also, this deletion could potentially affect the intracellular trafficking of ergosterol or drug molecules. Though the deletion of glutamic acid in the 98^th^ position is structurally disruptive, it enhances survival under antifungal pressure, representing a gain-of-function adaptation. Interestingly, 2 out of 3 mutations is present in all the lineages.

*AFR1* is one of the major ABC efflux pumps in *Cryptococcus* sp. that is specialised for azole extrusion. Though all the azoles disrupt sterol homeostasis, fluconazole’s weaker binding to CYP51, better efflux compatibility, chronic clinical usage and stronger stress-signalling induction make it the primary trigger for *AFR1* overexpression in *C. neoformans. AFR1* is a key player promoting adaptive resistance in *Cryptococcus* sp. AFR1 is present in almost all the study isolates, yet its expression is controlled by external factors.

*NRG1* is a transcription factor that regulates genes involved in cell wall organisation and integrity, capsule and melanin production. The absence of *NRG1* increases tolerance to azoles by upregulation of *AFR1* or by changing the sterol biosynthesis. Though *NRG1*’s role in promoting antifungal resistance is indirect, it is significant. In our study, isolates of only a few strains belonging to the VNB lineage show the absence of *NRG1*.

Altogether, for promoting azole resistance, upregulation of *AFR1* along with the point mutations in *ERG11*/ aneuploidy play a significant role.

The pyrimidine analogue resistance was conferred by mutations in *FUR1,* which is one of the most critical enzymes involved in the pyrimidine salvage pathway^29^. Followed by *FCY2* and *MSH2* and *NRG1*. The mutation/ absence of *MSH2* accelerates the acquisition of all the above mutations. *FCY2* encodes cytosine permease, a member of the major facilitator superfamily that imports 5-fluorocytosine into the cell. The mutation at the 157^th^ position, where the alanine (small, nonpolar) is replaced by valine (bulkier, hydrophobic), is a conservative substitution, but it alters the transport efficiency. This is a partial loss of function mutation that results in decreased intracellular drug, indirectly resisting by reducing drug entry. This *FCY2* mutation is present only in the VNIV strain of study isolates used. But rarely, literature also states that the VNI and VNII lineages also have the *FCY2* mutation. Mutations in the *FUR1* genes directly promote antifungal resistance by not allowing the 5-FC to convert into 5-FUMP, which is the crucial step to activate the drug. There are five *FUR1* mutations described to cause antifungal resistance and all those point mutations were non-conservative amino acid changes and result in disruptive or truncated protein. The mutation D42V causes protein destabilisation, resulting in disruption of the pyrimidine salvage pathway crucial for 5-FC activation. The mutation P140S introduces flexibility in the protein structure, resulting in a conformational shift, reducing *FUR1*’s catalytic activity, contributing to reduced activation of 5-FC, promoting moderate resistance. The mutation G87D has a high possibility of causing a loss-of-function mutation resulting in truncated proteins. The mutation G190E is not a part of the active site pocket but effectively disrupts the stability of the protein, indirectly hindering the enzyme activity. E64 is a conserved amino acid that plays an important role in protein stabilisation. The mutation of E64G results in protein destabilisation and hinders the pyrimidine salvage pathway. Among the five explained mutations, except D42V, all other mutations were widely present across all lineages of the study isolates. *MSH2* is a key component of the DNA mismatch repair system. The absence or mutations in *MSH2* lead to rapid acquisition of antifungal resistance, especially for azoles and pyrimidine analogues^28^. The major mutations in *MSH2* are K399E, N900D and D904G, which are all non-conservative substitutions leading to structural instability of the protein. The mutation position K399 was located in the core MMR domain that reduces mismatch repair recognition and repair efficacy. The mutation positions N900/ D904 are present in the C-terminal region, which is important for interaction with MSH6 or other repair proteins. The non-conservative mutations in these regions lead to complex destabilisation. The loss or reduced activity of this protein complex increases the mutation rate in genes *FUR1*, *FCY1* and *ERG11*. Also, this might lead to the emergence of hypermutator strains. In the study isolates, the C8 strain belonging to VNI, serotype A, has no *MSH2* gene and all the other strains have at least two of these three mutations in *MSH2,* suggesting higher chances of acquiring resistance to azoles and pyrimidine analogues. The mutations in *NRG1* suppress UDP-glucuronic acid accumulation, resulting in 5-FC resistance^40^.

## Conclusion

Comparative genomic study of 139 isolates of *C. neoformans* genomes collected across the continents between 1988 and 2021 culminates in key insights into the global genomic epidemiology of *C. neoformans*. The findings highlight the dominance of serotype A, the VNI lineage and ST5 worldwide. The pathogen exhibits low overall genetic diversity, with the highest variation found in Southern Africa and North America, suggesting S. Africa as a potential ancestral hotspot. The MATα mating type is widespread, while MATa is rare, indicating unisexual mating as a primary mechanism for reproduction and genetic diversity. Notably, the pangenome of *C. neoformans* is closed, which shows the stabilisation of essential functions and major virulence and antifungal resistance traits likely to reflect the species’ evolutionary history and adaptation to its niche. Yet it also shows enough accessory variations to permit lineage-specific adaptation and antifungal resistance. Although infection rates may be relatively low, likely due to niche-specific existence as basidiospores, the mortality rate for cryptococcosis remains high in immunocompromised individuals, especially in patients with advanced HIV, due to delayed diagnosis and limited treatment options.

## Supporting information

Supplementary table 1: Metadata from NCBI datasets along with the Serotype, molecular type, ST and Mating type

Supplementary table 2: Genome diversity profile - the table depicts the average nucleotide identity between the study genomes

Supplementary table 3: Whole genome SNP profile of study genomes (n=139)

Supplementary table 4: Pan-genome profile of study genomes (n=139)

Supplementary table 5: Major virulence genes (n=17) and their role in virulence

## Acknowledgment

We thank SASTRA Deemed University for providing the research facilities and infrastructure.

## Supplementary tables

**Supplementary Table 1:** Metadata from NCBI datasets along with the Serotype, molecular type, ST and Mating type.

**Supplementary Table 2:** Genome diversity profile - the table depicts the average nucleotide identity between the study genomes

**Supplementary Table 3:** Whole genome SNP profile of study genomes (n=139)

**Supplementary Table 4:** Pan-genome profile of study genomes (n=139)

**Supplementary Table 5:** Major virulence genes (n=17) and their role in virulence

