## Supplementary figures and images for "Deciphering the global genomic landscape of *C. neoformans*: Population dynamics, molecular epidemiology and genomic signatures of pathogenicity"

### Supplementary figure 1

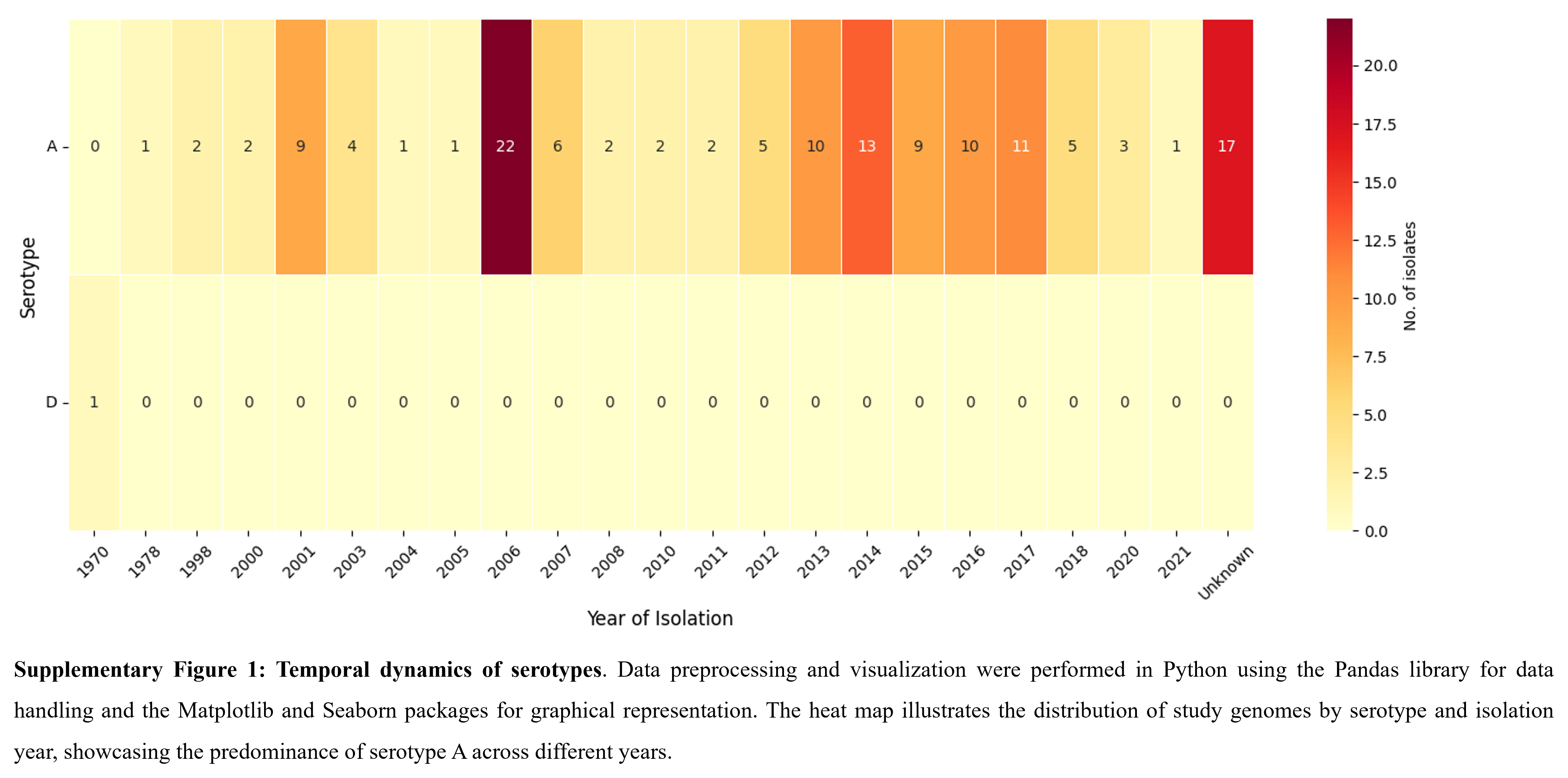

### Supplementary figure 2

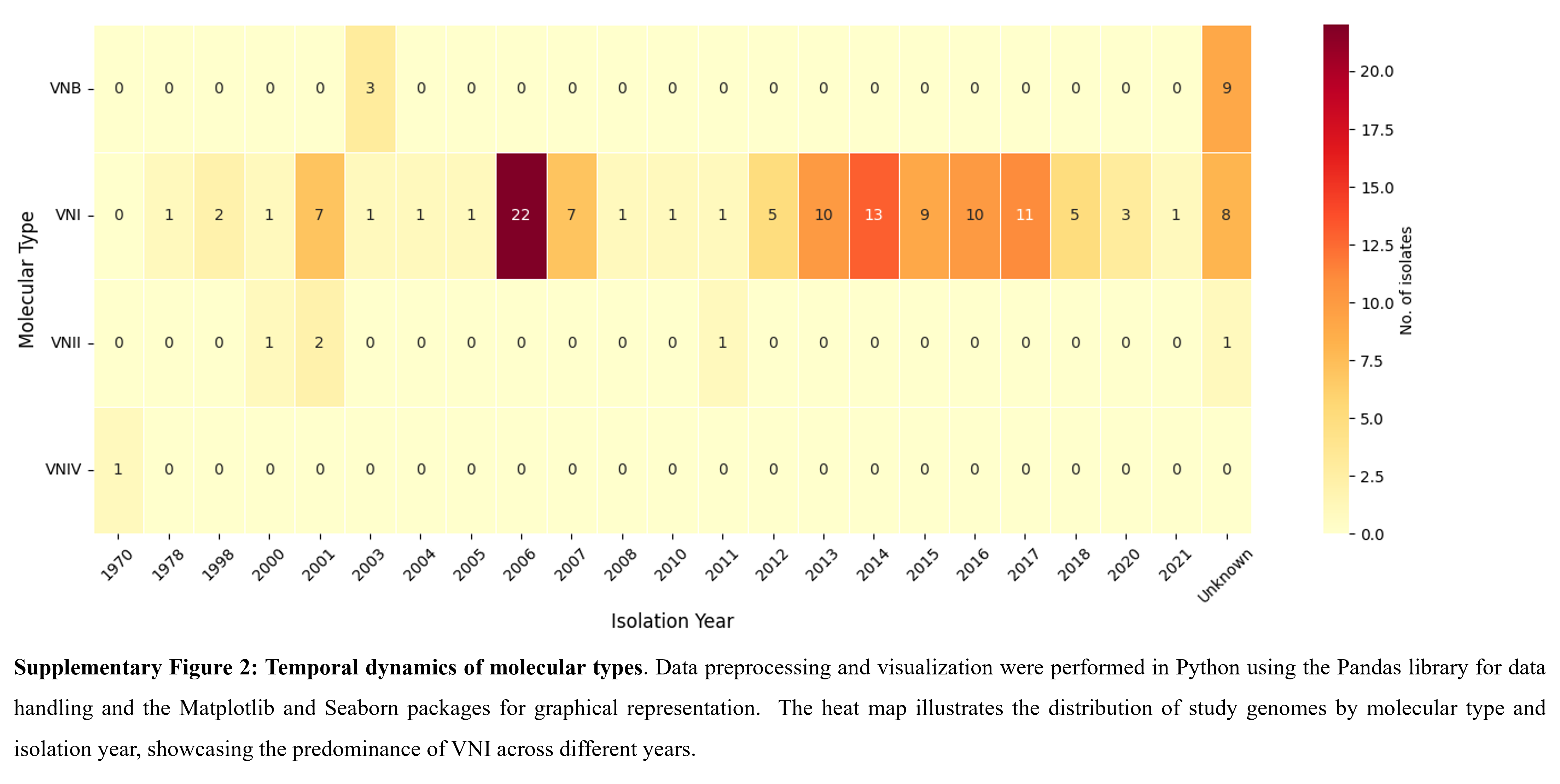

### Supplementary figure 3

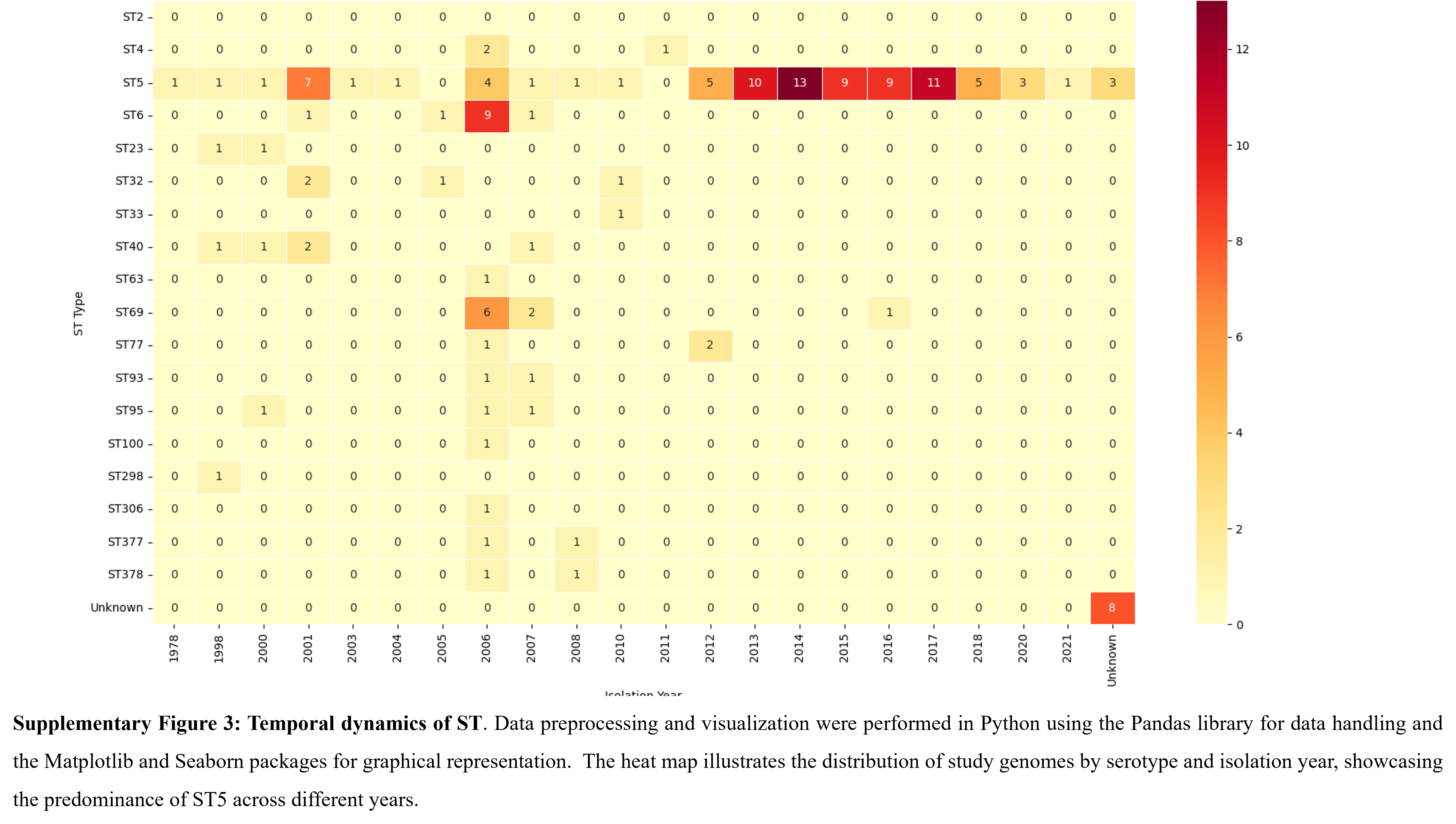

### Supplementary figure 4

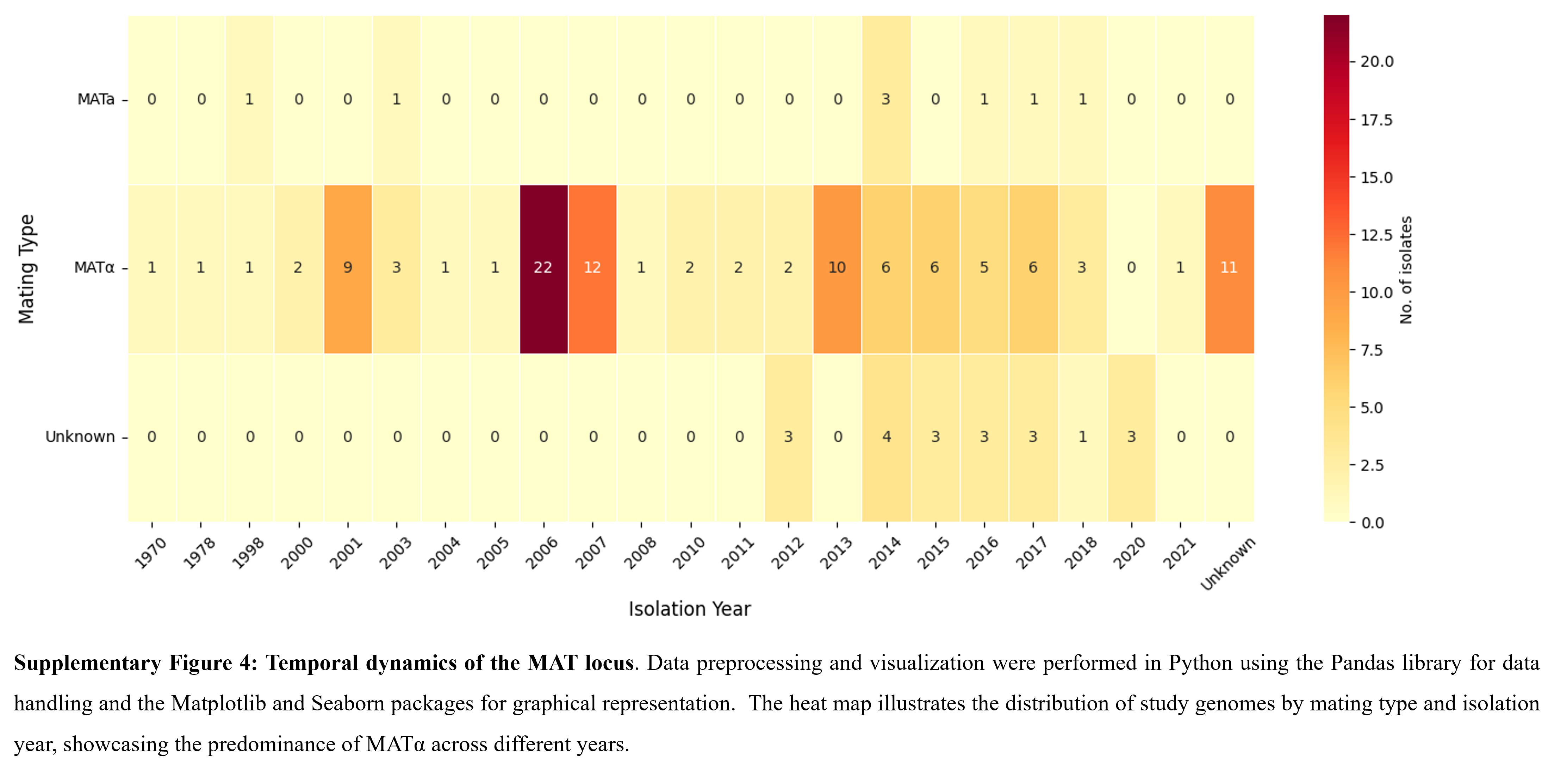
